# An acidophilic fungus is integral to prey digestion in a carnivorous plant

**DOI:** 10.1101/2023.11.07.566145

**Authors:** Pei-Feng Sun, Min R. Lu, Yu-Ching Liu, Yu-fei Lin, Daphne Z. Hoh, Huei-Mien Ke, I-Fan Wang, Mei-Yeh Jade Lu, Roland Kirschner, Ying-Chung Jimmy Lin, Ying-Lan Chen, Isheng Jason Tsai

## Abstract

Carnivorous plant leaves, such as those of the spoon-leaved sundew *Drosera spatulata*, secrete mucilage which hosts microorganisms potentially aiding in prey digestion. We characterised the mucilage microbial communities and identified the acidophilic fungus *Acrodontium crateriforme* as the ecologically dominant species. The fungus grows and sporulates on sundew glands as its preferred acidic environment. We show that the *A. crateriforme* has a reduced genome similar to that of other symbiotic fungi. Based on the transcriptomes when encountering prey insects, we revealed a high degree of genes co-option in each species during fungus-plant coexistence and digestion. Expression patterns of the holobiont during digestion further revealed synergistic effects in several gene families including fungal aspartic and sedolisin peptidases, facilitating the digestion of sundew’s prey, as well as transporters and dose-dependent responses in plant genes involved in jasmonate signalling pathway. This study establishes that botanical carnivory is defined by multidimensional adaptations correlated with interspecies interactions.

---

Botanical carnivory has evolved independently at least 11 times in the plant kingdom, each showcasing distinct unique molecular adaptations to attract, capture and digest insects^1^. Many carnivorous plants constitute iconic research models since the era of Charles Darwin^2^ to understand the evolutionary and molecular basis of their predatory abilities, which are frequently confined within a highly modified leaf organ. These specialised leaves secrete digestive exudates that contain a diverse array of microorganisms. Although the significance of microbiota in vertebrate digestion is widely established^3^, the symbiotic interplay between carnivorous plants and their associated microbiota is emerging in research^4^, and the underlying molecular responses to which microorganisms facilitate or enhance plant carnivory remain to be elucidated.

Plant-microbe interactions are highly dynamic and can impact plant fitness through many mechanisms^5^. Previous studies using metabarcoding suggested that the digestive mucilage encapsulated by the modified leaves, known as traps, is colonised by diverse communities of microorganisms (**Extended Data Table 1**). In bladderwort, corkscrew and pitcher plants, there were no dominant species present within the traps, but they can be broadly grouped into major bacterial phyla^6–11^. Bacterial diversity and biomass were found to improve prey decomposition rates in the pitcher plant *Darlingtonia californica* by increasing host leaves’ nitrogen uptake efficiency^11^. Meta-transcriptomic profiling of *Genlisea* species revealing non-host transcripts were dominated by metazoan hydrolases, suggesting a role in phosphate supplementation^12^. In addition, the composition of the microbiota appears to be highly time-dependent and influenced by factors such as host^11^, surrounding environment^13^, prey-associated bacteria^13^ and enhanced species dispersal capabilities^7^, highlighting the complex interplay of factors that shape these microbial communities.

Species of *Drosera*, known as sundews and part of the second largest carnivorous family after Lentibulariaceae^14^, have ‘flypaper’ leaves with tentacle-like trichomes^15^ that secrete mucilage to trap prey, with *D. spatulata* leaves enveloping the prey during digestion. After successful capture, the tentacles rapidly envelop the prey, which is then degraded and mineralised by the secreted digestive enzymes. This complex behaviour is stimulus specific, i.e., not activated by water droplets^16^, and is mediated by regulation of the jasmonate (JA) signalling pathway which was pre-established in non-carnivorous plants^16^. Recent sequencing and comparative genomics of carnivorous plants has revealed expansion and clade-specific gene families involved in carnivory. Members of these genes have been co-opted^17^ by acquiring new roles from ancestral processes ranging from defence, such as JA metabolism and signalling, to different stages of the capturing cycles including peptidases and hydrolases for prey digestion^18^. The extent to which these genes still retain their ancestral functions remains to be elucidated. Another outstanding question is whether the outcome of the inter-species interactions also drives genomic and transcriptomic adaptations to the microbes in the holobiont^19^.

To investigate the potential inter-species interactions, we focus on *Drosera spatulata*^20^ (**Fig. 1a**), a sundew native to tropical regions including Taiwan. The fascinating mechanism of trap movement in sundews, associated with prey digestion, has long attracted scientific curiosity due to its complex operational intricacies^21^, but importantly, act as a tractable response to diverse stimuli^22^ in both natural and laboratory settings. This species, with its sequenced genome^18^ and documented microbial presence on its phyllosphere — where surface yeast has been shown to promote growth^23^ — collectively present a model microbial ecosystem to experimentally interrogate the underlying molecular details. We sought to characterise the mucilage microbial community and assess their impact on digestion. By exploring these aspects, our research aims to shed light on the intricate symbiotic interactions between carnivorous plants and their resident microbes.

**Fig. 1.**
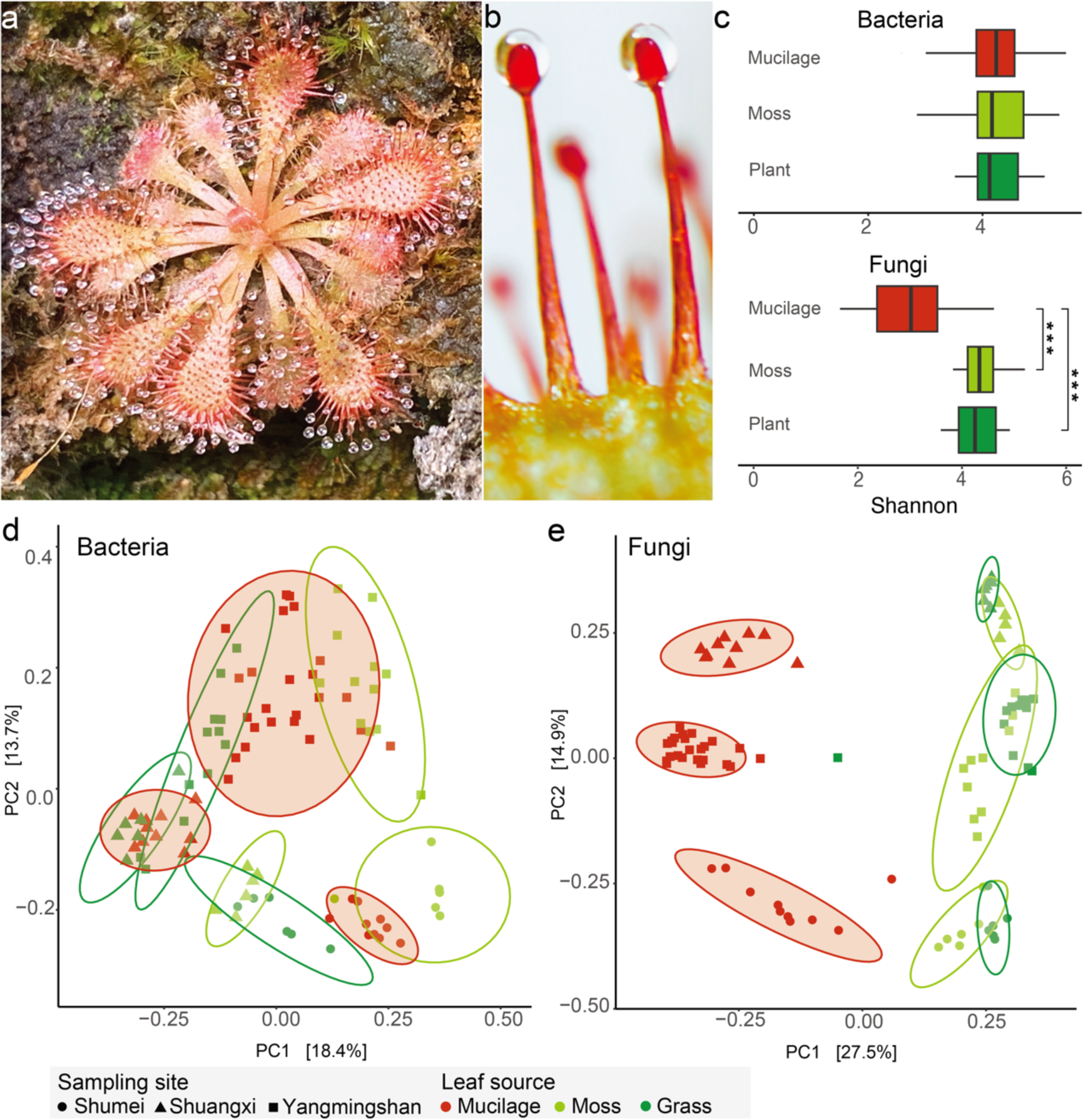
Microbial Communities of *Drosera spatulata* mucilage and surrounding environment. **a.** *Drosera spatulata* and **b.** close up of stalk glands with secreted mucilage. **c.** Bacterial and fungal species evenness in sundew mucilage versus neighbouring plant leaf surfaces. Asterisk denote significant difference from Wilcoxon rank sum test (*** indicate P<0.001). Beta diversity (Bray-Curtis index) of **d.** bacterial and **e.** fungal communities. Eclipses were drawn at 95% confidence level within samples of same source and site.

## Results

### Fungal communities in *Drosera* mucilage were different to surrounding environment

To better understand the microbial diversity and composition of the mucilage found on the sundew *Drosera spatulata*, we employed sterilised filter papers to collect mucilage from sundews (**Fig. 1b)** and surrounding plants typically found on cliff habitats of Northern Taiwan (**Supplementary Fig. 1**). A total of 92 samples were subjected to 16S and ITS amplicon metabarcoding (**Supplementary Table 1**). Each sample had average of 546 and 445 bacterial and fungal operational taxonomic units (OTUs), respectively. The bacterial species evenness was comparable between the sundew mucilage and leaf surfaces of surrounding plants (**Fig. 1c**, Wilcoxon rank sum test, P=0.81). In contrast, the fungal communities in mucilage had a significantly reduced species evenness (**Fig. 1c**). Beta diversity of microbial communities estimated using the Bray-Curtis index indicated no discernible differences in bacterial communities between leaf surfaces (**Fig. 1d**, PERMANOVA, Leaf surface: R^2^=0.12, P=0.06). 42.9% of OTUs corresponding to 53.8-99.9% relative abundance of mucilage microbiome were also found in the adjacent samples (**Supplementary Table 2**), implying that the mucilage bacterial community resembled that of the surrounding environments. However, the mucilage fungal community was significant different compared to surrounding samples (**Fig. 1e**, PERMANOVA, Leaf surface: R^2^=0.30, P=0.001). These results suggest the presence of a dominant fungal species in the *D. spatulata* mucilage.

### Ecological dominance of *A. crateriforme* in *D. spatulata* mucilage

We examined the relative abundances of the top five major bacterial and fungal taxa among leaf surfaces and revealed a single dominant fungal operational taxonomic unit (OTU) in the mucilage samples averaging 46.6% relative abundance (**Fig. 2a**). Reanalysis of beta diversity excluding this OTU have rendered the fungal communities with no significant differences between leaf hosts (**Supplementary Fig. 2;** PERMANOVA, R^2^=0.1197, P=0.06 suggesting that this OTU was the main biotic factor affecting the fungal community structures.

**Fig. 2.**
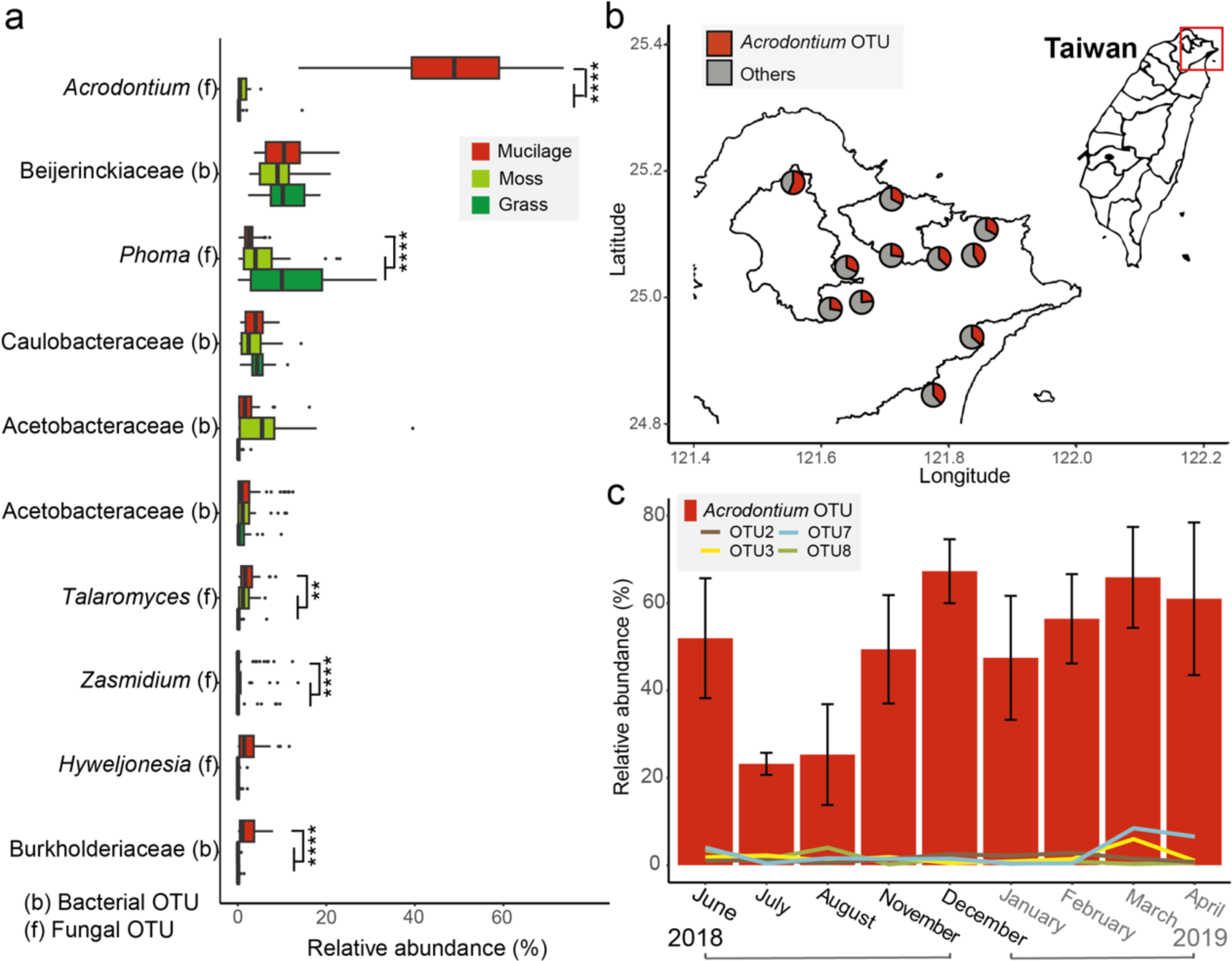
Ecologically dominance of *Acrodontium* OTU in *D. spatulata* mucilage. **a.** Relative abundances of the top ten dominant bacterial [marked as (b)] or fungal [marked as (f)] taxa among leaf surfaces, showcasing the *Acrodontium* OTU as the dominant species in the mucilage. **b.** Relative abundance of *Acrodontium* in 52 additional sundew mucilage samples across Northern Taiwan. Red colour in pie chart denotes relative abundance of *Acrodontium* OTU **c.** Temporal variation of *Acrodontium* and the next four major OTUs’ relative abundance in mucilage over nine months between 2018-2019 at two sites.

The taxonomic placement of the dominant fungal OTU belonged to the genus *Acrodontium*. By re-analysing the available information from Globalfungi^24^ database, it was revealed 1.2% of 57,184 samples harboured *Acrodontium* from forests (77.3%), followed by grasslands (12.2%), with the majority of relative abundances being less than 1% (**Supplementary Fig. 3 and Supplementary Table 3**). *Acrodontium* appeared to be preferentially located in more acidic samples (**Supplementary Fig. 4**), which is considered an ancestral trait as it is phylogenetically closely related to a group of fungi that can thrive at low pH^25^. To investigate the extent of *Acrodontium* dominance in *D. spatulata*, additional sequencing of mucilage samples spanning ∼45 km of Northern Taiwan revealed its remarkable dominance in all samples with an average relative abundance of 30.8% (**Fig. 2b**). Continuous sampling of mucilage across two sites for nine months revealed that *Acrodontium* maintained its status as the most dominant fungus species despite a reduced 15.7% relative abundance during July and August (**Fig. 2c**). This further validates the ecologically dominance of *Acrodontium* in the mucilage of *D. spatulata* irrespective of spatial and temporal factors.

To investigate the roles of fungi to the carnivorous plant host, we isolated the two most dominant OTUs from fresh mucilage of *D. spatulata* and identified them as *Acrodontium crateriforme* and *Phoma herbarum* of the families Teratosphaeriaceae and Didymellaceae, respectively (**Supplementary Fig. 5 and 6**). The latter is a plant pathogen causing leaf spot in various crops^26^, while *A. crateriforme* is cosmopolitan in nature found in soil^27^, plant material^28,29^, compost^30^, air^31^ and rock surfaces^32^. Interestingly, *A. crateriforme* was isolated in the digestive fluid of the Indian pitcher plant *Nepenthes khasiana*^33^ and preferred nitrogen-rich substrates^34^. We reanalysed 14 published fungal metabarcoding datasets of carnivorous plants, and detected its presence in purple pitcher plant *Sarracenia purpurea*^35^. Despite having a lower relative abundance of 0.2-1.4% (**Extended Data Table 1**), there was a general presence of *A. crateriforme* in carnivorous plants.

Like *D. spatulata, A. crateriforme* is well suited to grow under laboratory conditions, and can be cultivated using PDA and MS media. *A. crateriforme* grows optimally at pH 4–5 mirroring the acidity of *D. spatulata* mucilage^36–38^ (**Supplementary Fig. 7**), suggesting its acidophilic nature. In contrast, *Ph. herbarum* preferred a more neutral pH. The optimal culture temperature for *A. crateriforme* is 25°C (**Supplementary Fig. 8**), aligning with *D. spatulata’s* growth range (7‒32°C) and the average monthly temperature of 22.0°C at the sampling sites (**Supplementary Fig. 9**). The summer temperature peaks (29– 35°C) at the sites may, combined with biotic factors such as optimal growth of *Ph. herbarum* at this higher temperature range (**Supplementary Fig. 8**), explain the reduced abundance of *A. crateriforme* at higher temperature (**Fig. 2c**).

### Plant-fungus coexistence increases digestive performance of sundews

Examining the stalk glands of *D. spatulata* growing in a sterilised environment using a scanning electron microscope (SEM) revealed clear surfaces (**Fig. 3a and Extended Data Fig. 1a-c**). Inoculating the sundew with *A. crateriforme* revealed hyphae that grew over the glands (**Fig. 3b and Extended Data Fig. 1d-f**), while conidiophores and detached conidia were observed in glands collected from the wild (**Fig. 3c**). This observation demonstrates that *A. crateriforme* colonises and reproduces on the sundew stalk glands^39^, and the mucilage harbours free hyphae or conidiophores. The cultivation of *A. crateriforme* is positively correlated with the amount of *Polyrhachis dives* ant powder added to the medium, suggesting that the fungus can utilise insects as a growth supplement (**Supplementary Fig. 10**).

**Fig. 3.**
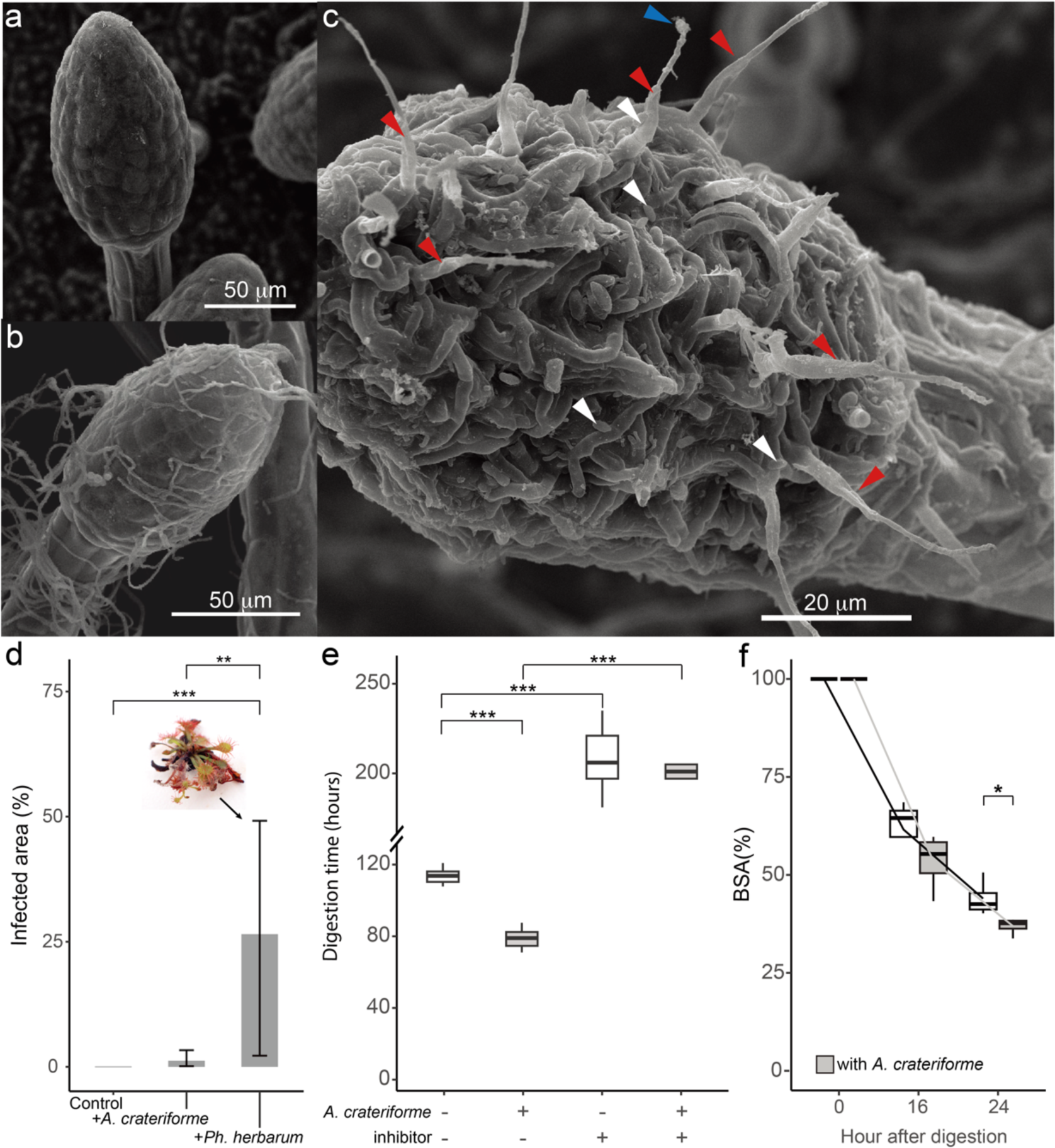
The *A. crateriforme*-*Drosera spatulata* holobiont. Scanning electron microscope (SEM) image of sundew stalk glands under **a.** sterilised conditions, **b.** inoculated with *A. crateriforme*, and **c.** natural habitat. Different arrow colour denote conidiophores from which conidia were already detached (red), a conidium attached to the tip of a conidiophore (blue), and detached conidia (white). **d.** Effects on *D. spatulata* post-inoculation with *A. crateriforme* and *Ph. herbarum*. A photo showing wilt of *D. spatulata* as a result of *Ph. herbarum*. **e.** Re-opening time of sundew traps supplementing with ant powder in different treatments. **f.** Application of biotin-labelled BSA during sundew digestion. Asterisk denote P values from Wilcoxon-rank sum test (* P<0.05, ** P<0.01, *** P<0.001). + and – denote presence and absence of treatment, respectively.

To establish the potential contribution of *A. crateriforme* to the sundew host, *A. crateriforme* was first inoculated onto the *D. spatulata* leaves and there were no significant differences in the plant morphology and net weight after one month (**Fig. 3d** and **Supplementary Fig. 11**). Conversely, inoculating *Ph. herbarum* with the same concentration resulted in plant wilt (**Fig. 3d**), suggesting stable plant-fungus coexistence between *A. crateriforme* and *D. spatulata*. When supplementing leaves with ant powder, the times when the stalk glands fully covered the prey and when they returned to their original position were recorded. The period between these two points of time indicates the reopening of the trap and completion of digestion. The inoculated samples’ trap re-opening was 26.4% earlier than in non-inoculated samples (averaging 106.2 versus 78.1 hours; **Fig. 3e**), demonstrating the fungus’s direct involvement in sundew digestion. The addition of protease inhibitor significantly lengthened the reopening time in both treatments, corroborating previous findings that the peptidases were primarily involved during the digestion process^40–42^. Application of biotinylated bovine serum albumin on the collected mucilage showed a declining trend during digestion and was significantly reduced after 24 hours from the *A. crateriforme* inoculated-samples (inoculated sample: 36.9 % vs without: 44.0%, P=0.03, Wilcoxon rank sum exact test, **Fig. 3f and Supplementary Fig. 12**), emphasising that more proteins were being digested during this time with the presence of *A. crateriforme*. Together, the results suggest that *A. crateriforme* is part of the sundew holobiont and enhanced the digestion process.

### Genome of *A. crateriforme* as an extremophilic fungus

We sequenced and assembled the *A. crateriforme* genome using 10.5Gb of Oxford Nanopore long reads and polished the consensus sequences with Illumina reads. The final assembly resulted in 14 contigs, with 13 containing TTAGGG copies at both ends corresponding to gapless chromosomes (**Supplementary Table 4**). The assembly size of 23.1 Mb represented the first genome from the genus *Acrodontium*. We predicted 8,030 gene models using MAKER pipeline^43^ aided by RNAseq as hints. Of these, 97.3% of the predicted gene models were found to be orthologous to at least one of the 25 representative species in the order Capnodiales (**Supplementary Table 5**), suggesting a conserved core genome with potential unique adaptations. A species phylogeny was constructed by coalescencing 9,757 orthogroup trees, which placed *A. crateriforme* within a group of extremophilic species (**Fig. 4a**) including well-known acidophiles such as *Acidomyces richnondensis* and *Hortaea acidophila*^44^.

**Fig. 4.**
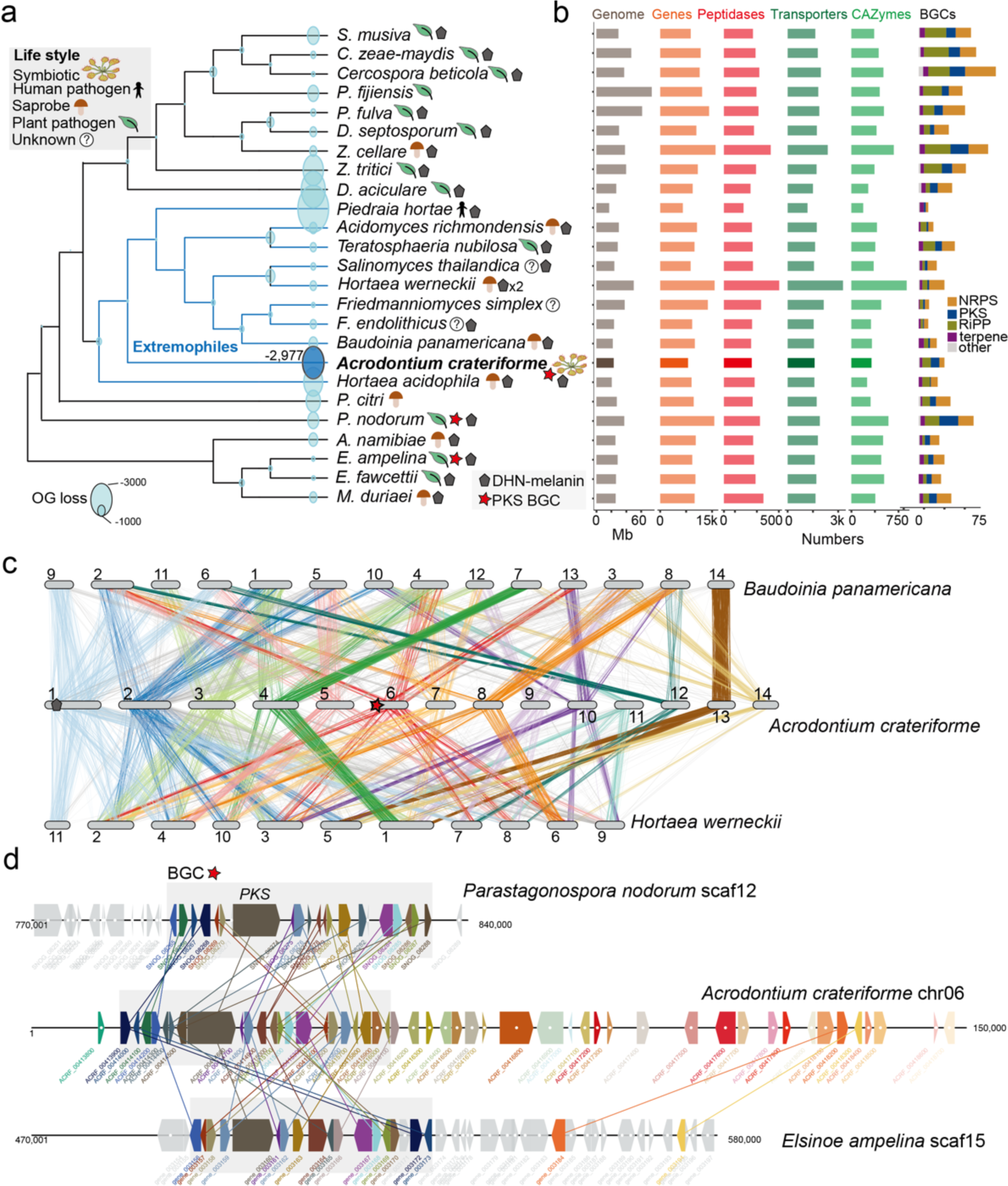
Genomic features of *A. crateriforme*. **a.** Phylogenetic placement of *A. crateriforme* among extremophilic fungi denoted in blue branches, highlighting its association with known acidophiles. All nodes have a 100% bootstrap support. Number next to *A. crateriforme* denote number of lost OGs inferred by DOLLOP. **b.** Genome description and functional annotations of the *A. crateriforme* proteome. **c.** Chromosomal rearrangements amongst extremophiles through clustering of single copy ortholog pairs. Colour designate *A. crateriforme* **d.** Synteny between a subtelomeric polyketide cluster on *A. crateriforme* chromosome six and plant pathogens *E. ampelina* and *P. nodorum*. *A. crateriforme* genes and orthologs were coloured sequentially.

We functionally annotated the *A. crateriforme* proteome to identify genes and gene families associated to its ecology and metabolism (**Fig. 4b**). PCA of protein family domain numbers from each species first differentiated the extremophiles *Friedmanniomyces simplex* and *Hortaea werneckii* from others with their partial^45^ or whole^46^ duplicated genome (**Supplementary Fig. 13a**). *A. crateriforme* was positioned between its extremophile relatives and outgroup plant pathogens (**Supplementary Fig. 13b**), suggesting that gene family dynamics were influenced by adaptation to an acidic environment as well as plant association. Although *A. crateriforme* contained relatively fewer gene models, it was not a reflection of assembly quality, with 98.9% completeness based on BUSCO^47,48^ (**Supplementary Table 5**) and encodes a similar number of CAZymes, peptidases, transporters and BGCs to its sister species (**Fig. 4b**). Amongst the extremophiles, inference of gene family dynamics indicated a relatively high number of losses in *A. crateriforme* compared to the human dermatophyte *Piedraia hortae* (**Fig. 4a**). *A. crateriforme* have lost members of glycoside hydrolase 6, 11, 28, and 43 which degrade plant cell wall and their losses have been implicated as signatures of symbiotic fungi^49^ such as ectomycorrhizal fungi^50^ (**Supplementary Fig. 14 and Supplementary Table 6**). Further specialisations of *A. crateriforme* include a higher number of polyketide synthase clusters within this group of species (**Fig. 4b**) Most of the identified BGCs in *A. crateriforme* were unique and not shared with other representative species, implying its distinct profile of secondary metabolites, particularly the polyketides (**Supplementary Fig. 15**). Interestingly, *A. crateriforme* encodes two Neprosin domain-containing genes (**Supplementary Fig. 16**), which were absent in all representative species and rare in fungi (149 versus 8,118 in Viridiplantae; InterPro, last assessed October 2023). Neprosin was first discovered in Raffles’ pitcher plant *Nepenthes rafflesiana* as a novel peptidase capable of digesting proteins at low concentrations without substrate size restriction^51^, hinting at its potential involvement in prey digestion.

The role of genomic rearrangements, especially in subtelomeric regions, in fungal evolution has been well-documented, particularly in pathogenic species ^52,53^. We sought to characterise the mode of genome evolution in this group of extremophilic fungi, and identified on average 4,747 pairwise single copy orthologs between *A. crateriforme* and sister extremophiles. Clustering of these orthologs with corresponding *A. crateriforme* chromosomes identified only one one-to-one linkage group of chromosome 13 (**Fig. 4c**), suggesting frequent chromosomal fusions and fissions since their last common ancestor. Gene order within linkage groups has been lost (**Supplementary Fig. 17**), suggesting extensive intra-chromosomal rearrangements, which appear to be a hallmark of genome evolution in the Capnodiales^54^. Such high genomic plasticity often led to the high turnover of gene family dynamics or emergence of biosynthetic gene clusters (BGCs) capable of producing novel secondary metabolites^55^. In the case of *A. crateriforme*, BGCs were enriched at subtelomeres (12/26 in subtelomeres with Observed to Expected ratio of 4.7; **Supplementary Fig. 18**). We identified a case of one polyketide cluster located on the end of chromosome six, which is shared with the plant pathogens *Elsinoe ampelina* and *Parastagonospora nodorum* (**Fig. 4d**), presumably as a result of *A. crateriforme* constantly encountering a plant-associated environment.

### Digestion-related genes were co-opted and retained ancestral expression trends from plant-microbial coexistence

To dissect how sundew and *A. crateriforme* respond to each other or encountering insect at the transcriptome level, we first generated gene expression data from both species cultured under minimal nutrient conditions. This baseline data was then compared to two scenarios: application to ant powder indicative of digestion and fungal inoculation onto sundew leaves denoting coexistence (**Supplementary Fig. 19**). Remarkably, 61.3–63.9% of differentially expressed genes (DEGs) identified during the digestion phase for each species were co-expressed in the coexistence phase (**Fig. 5a**), suggesting an intrinsic regulatory synergy between the two processes. Gene ontology (GO) analysis revealed that more than half of the GO term enrichment of the up-regulated genes overlapped between the two conditions in *D. spatulata,* with the most significant terms including secondary metabolic process, response to chemical and other organism (**Supplementary Fig. 20 and Supplementary Table 7**). This suggests that the majority of plant genes that were involved in defence mechanisms^56,57^ have been co-opted in the digestion process but still retained their ancestral functions. An example include members of plant chitinase (GH18 and GH19) (**Supplementary Fig. 21**), which have roles in digestion and original functions in defence against phytopathogen^58,59^. However, this degree of overlap was not observed in *A. crateriforme* (**Supplementary Table 8**), reflecting independent specialisations between the two species. The coexistence phase yielded more DEGs compared to the digestion phase (**Fig. 5a)**, consistent with that the former process being ancestral^60^. Interestingly, despite showing increased expression in both digestion and coexistence phases, the *D. spatulata* ammonium transporters which are central to NH_4_^+^ uptake during digestion^61^ exhibited a higher expression in the latter (**Supplementary Fig. 22**). The *A. crateriforme* ammonium transporters as well as nitrate reductase also exhibited the same expression trends (**Supplementary Fig. 22**), suggesting active ammonium exchange and utilisation already taking place within the plant-fungus holobiont.

**Fig 5.**
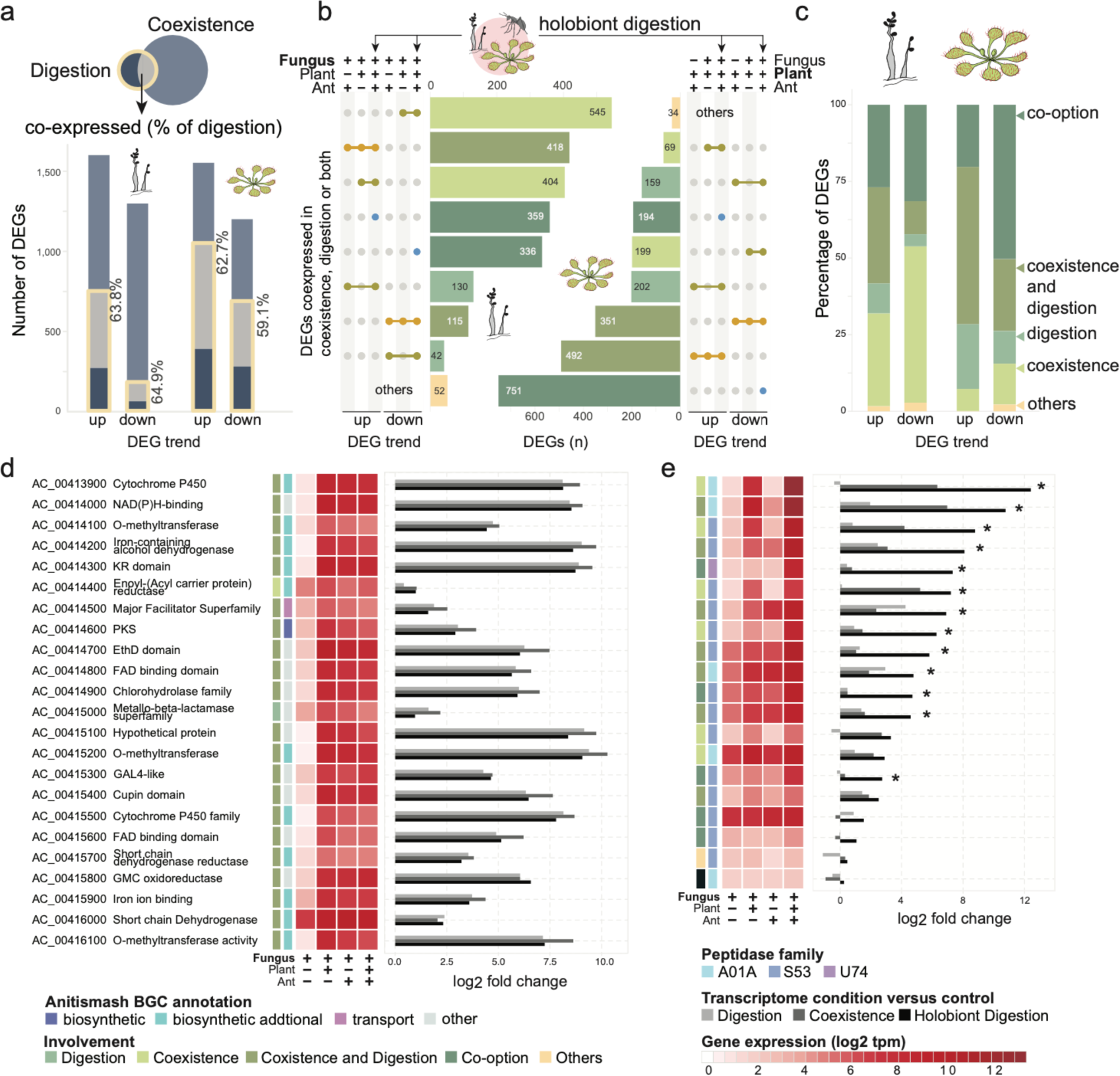
Transcriptome of the *D. spatulata*-*A. crateriforme* holobiont during digestion. **a.** Overlap of DEGs during digestion and coexistence for both species. **b.** Transcriptome profiling of the plant-fungus holobiont during digestion. DEGs were compared whether the same trends were observed in either digestion or coexistence process. **c.** Schematic representation of the role of the DEGs involved in the holobiont digestion. **d.** Upregulation of a BGC on chromosome six in *A. crateriforme*. **e.** Expression of representative fungal peptidases in a co-expression module (module 2 in **Supplementary** Figure 23) showing synergistic effects when both plant and insect prey are present. Asterisks indicate significantly upregulated expression determined by DEseq2 (adjusted P<0.05) between holobiont digestion and either digestion or coexistence phase.

### Transcriptome dynamic of holobiont digestion in nature

In nature, the digestion of insects took place in the mucilage of *D. spatulata*, with arthropod remains adhering to the stalk glands, where *A. crateriforme* was observed growing over the insect surface (**Extended Data Fig. 2**). To elucidate the mechanisms of carnivorous holobiont digestion in nature, we further characterised the holobiont transcriptome when supplemented with ant powder (**Supplementary Fig. 19**). We identified 3,554 and 3,662 DEGs in *A. crateriforme* and *D. spatulata*, respectively, with the majority of these genes exhibiting the same expression trend in at least one of the digestion or coexistence phases (**Fig. 5b**). These observations led us to propose a scenario in which the DEGs were concerned with one or both processes. In addition, we defined a new gene-option category in which the genes were only significantly differentially expressed when stimuli from both the interacting partner and the exogenous nutrient were present (**Fig. 5c**). More than half of upregulated DEGs in both species were designated having multiple or gene-opted, suggesting co-evolution and optimisation of the plant holobiont transcriptome as a result of constantly encountering each other and insect prey^19^. The *D. spatulata* DEGs had a higher proportion of multi-function and additive DEGs than the fungus, consistent with the optimisation of the genes repurposed to involve digestion^18^ while the majority still retained the ancestral function of species interaction.

Within *A. crateriforme*, the highest number of DEGs were categorised as involved in coexistence, suggesting its primary role in species interaction. Nevertheless, 21.1% of fungal DEGs were multi-functional. An example of this is the aforementioned BGC on chromosome six (**Fig. 4d**), which showed a consistent upregulation across all three phases (**Fig. 5d**) highlighting the need to effectively respond to multiple stimuli in natural environments. Interestingly, GO term enrichment of condition-specific genes revealed an opposite trend of up- and down-regulation of genes involved in the fungal and plant cell cycle, respectively (**Supplementary Table 9**), suggesting divergent responses in both species when faced with similar environment.

### Synergistic expression in fungal peptidases and transporters

Insect digestion in carnivorous plants is a well-coordinated process, first by synthesis and secretion of digestive enzymes to break down nutrients, followed by assimilation of nutrients with specialised transporters^61,62^. To investigate the putative roles of digestion in these gene families, additional transcriptome sequencing was performed in *D. spatulata* towards the end of digestion process (**Methods**). The regulation of peptidases during the different phases was determined using weighted correlation network analysis (WGCNA) ^63^, which identified five and nine co-expression modules in *A.* Genomic features of *A. crateriforme* Genomic features of *A. crateriforme* (**Supplementary Fig. 23 and 24**). The most dominant secreted peptidases in *D. spatulata* belonged to the cysteine (MEROPS^64^: C1) and aspartic families (MEROPS: A1). Droserasin, which has been implicated in digestion^22,40^, was contained in two co-expression modules (**Supplementary Fig. 24**). The two modules differed in that one module include genes that were constitutively up-regulated across conditions, whereas the other module contained genes that were only up-regulated during digestion by its own. In contrast, most of the highly expressed peptidases in *A. crateriforme* belonged to a co-expression module harbouring two copies of aspartic peptidase (AC_00151800 and AC_00417900), the entire sedolisin^65^ family (14/14 copies; MEROPS: S53) associated with increasing acidity and plant-associated lifestyle^66^, and a fungal copy with the aforementioned Neprosin domain (**Supplementary Fig. 23**). These genes demonstrated a synergistic effect in expression in prey digestion during co-existence, for instance the aspartic peptidases emerged as the dominant and third dominant entities across the entire transcriptome, with their expression levels increasing up to sixfold compared to either condition (**Fig. 5e**). The same trend was also observed in potassium, amino acid, oligopeptide and sugar transporters (**Supplementary Fig. 25**). Taken together, the results indicated *A. crateriforme*’s potential role in facilitating and benefiting from digestion in response to the combined signal of host and nutrient.

### Dosage dependent response of genes involved in Jasmonate (JA) signalling pathway

How carnivorous plant respond specifically either to symbiotic microbe or prey can be challenging, as the phytohormones are accumulated to elicit pathogen resistance in plants which induce genes that were also involved in digestion to prey stimuli^22,56^. We quantified the changes in *D. spatulata* phytohormone level after treatment with insect or different fungi. The application of both ant powder and microbes on the leaves have significantly increased the amount of Jasmonoyl-L-isoleucine (JA-Ile), a bioactive molecule of JA, after two hours (**Fig. 6a**) representing early response. In contrast, differences in salicylic acid and abscisic acid between control and treatments were not observed (**Supplementary Fig. 26**). The presence of *A. crateriforme* has increased JA-lle at the level between ant prey and the plant pathogen *Ph. herbarum*, suggesting a dose-dependent effect of JA in response to different biotic stimuli. We found that genes involved in JA signalling pathways were co-expressed in both coexistence and digestion phases (**Fig. 6b**), with the latter exhibiting heightened expressions as a late response towards the end of digestion process.

**Figure 6:**
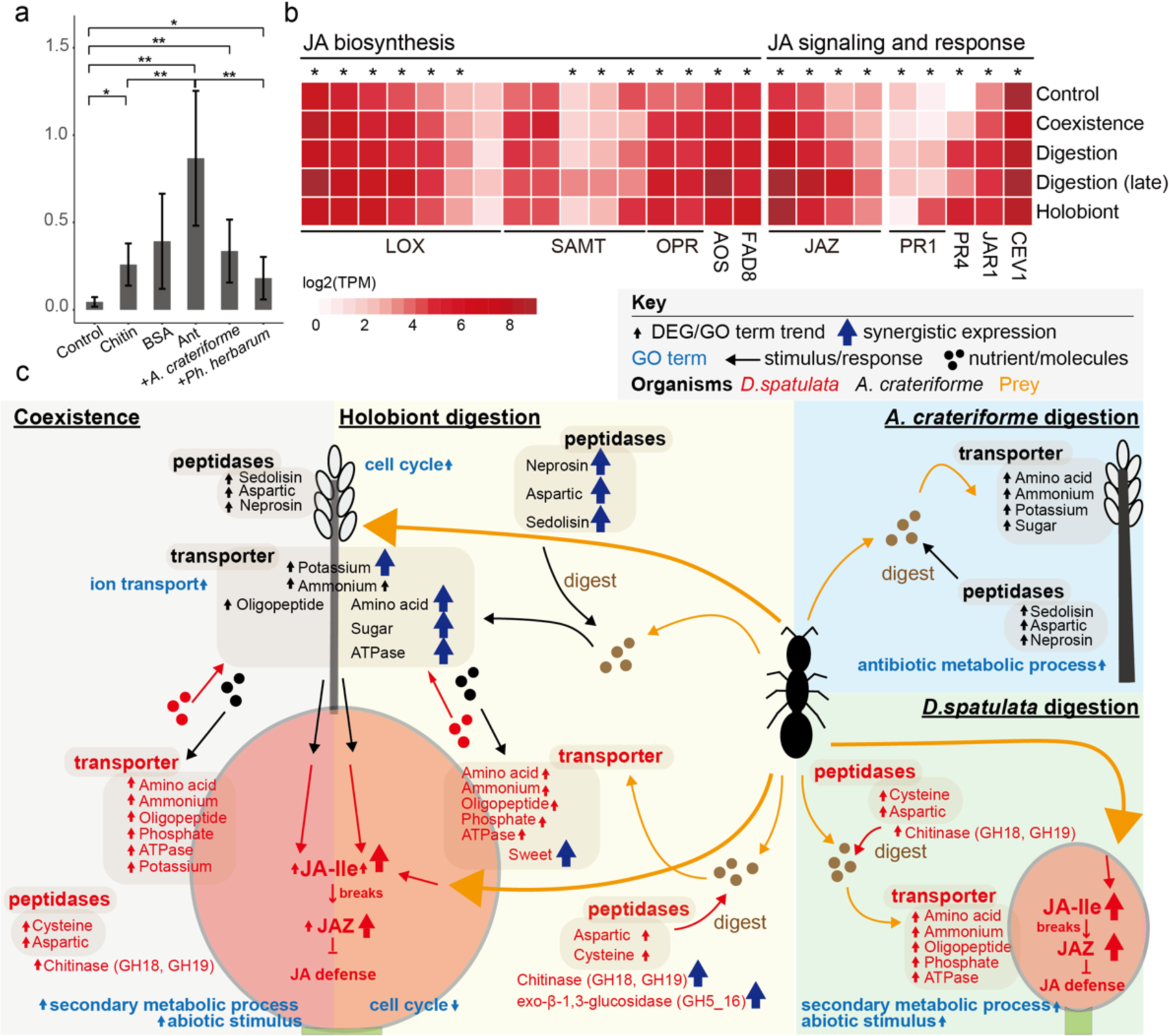
Interactions within the *D. spatulata*-*A. crateriforme* holobiont. **a.** Accumulation of Jasmonoyl-L-isoleucine (JA-Ile) levels in *D. spatulata* following various treatments. (Wilcoxon rank sum test; * P<0.05. ** P<0.01, *** P<0.001) **b.** Expression of genes involved in the JA signalling pathway during different phases. Asterisk denote genes that exhibited highest expression in the digestion phase. **c.** Holobiont response with each other and during digestion. A schematic diagram of *D. spatulata* stalk gland and *A. crateriforme* conidiophore with a summary of gene expression changes identified in this study are shown. Genes that are co-expressed in different phases are shown multiple times.

## Discussion

In this study, we elucidate a definitive symbiotic interaction between the carnivorous sundew *Drosera spatulata* and the acidophilic fungus *A. crateriforme* (**Fig. 6c**), providing insights that reshape views of plant versus prey in botanical carnivory since Darwin’s foundational work^2^. We show that the digestion time of insects by sundews was reduced by almost a quarter in the presence of the fungus, and that such cooperation is likely rooted in the common adaptative challenges that both carnivorous plants and extremophilic fungi face in harsh environments with minimal nutrients. This dynamic plant-fungus interplay reveals a multi-dimensional adaptation that goes beyond the conventional understanding of botanical carnivory.

The ecological dominance of *A. crateriforme* in sundews represents a rare case of intimate plant-fungus coexistence besides plant parasitism, as foliar microbiota are strongly influenced by both abiotic and biotic factors including leaf area, niche variability and the available resources^13^. The restrictive nature of *D. spatulata* mucilage’s acidity could potentially constrain its microbial interaction spectrum, a phenomenon echoed by observations of selective microbial compositions in adverse environments^67^. As an acidophilic species, resisting this biotic selection may be a first step in establishing the relationship. The dominance of a fungus in the *Drosera* microbiome that was not observed in other carnivorous plants may be correlated with constant drought stress by air-exposure of the stalked glands in contrast to the greater extent liquid medium contained in the traps of other carnivorous plants. Once colonising the leaf glands, *A. crateriforme* underwent a series of genomic changes to cope with the symbiotic life style and may increase the number of preys captured by reducing digestion time, implying that the coexistence is cooperative and may be mutualistic, as the level of prey capture is positively associated with plant fitness^68^ and may be more relevant as carnivorous plants are considered sit-and-wait predators. Considering that nutrient uptake efficiency in carnivorous plants is usually low, for example nitrogen uptake was less than half (29–42%)^69^, it is quite likely that *A. crateriforme* is able to utilise the unspent prey nutrients together with other microbial members. As fungi act as mediators between host and ecosystems^70^, it remains to be seen whether *A. crateriforme* will exhibit a context-dependent trophic level during its life cycle in *D. spatulata* and other carnivorous plant^33^. Inter-species interactions will be an important component in future cost-benefit models to explain how carnivorous plants survive the harsh habitat^71^.

Although it is generally accepted that plant carnivory genes are evolved from defence mechanisms, the availability of symbiotic *A. crateriforme* and single or dual-species transcriptomic landscape during the digestion phase enabled us to delineate the relative contribution of fungus and plant as well as the role of each gene. The extent of co-opted genes appeared more pronounced in *D. spatulata*, many of which maintained their expression trends in both digestion and coexistence phases. This suggests that *D. spatulata* employs the same set of genes to mediate defence in response to biotic stimuli from microorganisms and prioritises over digestion in the presence of prey which may be regulated by JA pathways. From a microbe’s perspective, the proteome of *A. crateriforme* also shows gene co-option as well as synergistic expression suggesting that the fungus has co-evolved with the sundew host to actively facilitate both processes in the shared environment^19^. For example, fungal sedolisins have been associated with increasing acidity and a plant-associated lifestyle^66^. The collective peptidases of both species can be utilised to degrade large proteins/peptides on acidic mucilage to generate nutrients in decomposing insect prey, potentially increasing overall digested nutrient levels.

In summary, our results provide direct evidence that microorganisms are integral to prey digestion in carnivorous plants (**Fig. 6c**). Our work supports the idea that plant-microbial interactions have been selected during evolution to increase the overall fitness of holobiont^19^. *Drosera-Acrodontium* is an amenable laboratory system, since both can be grown separately and together in the laboratory. We hypothesise just like plant carnivory has been independently evolved with convergence in different plant groups, plant-microbial interactions capable of facilitating the process digestion are likely to emerge in different carnivorous plants. Microbial ecosystems in other carnivorous plants can be highly complex that contain predators of microorganisms amongst inquilines^72^, and we provide an initial framework for detangling these relationships.

## Methods and materials

More detailed information on the materials and methods used in this study are provided in **Supplementary Information**.

### Genomic DNA extraction and metabarcoding of environmental samples

Details of sampling are provided in the **Supplementary Information**. Total genomic DNA was extracted from filter papers using modified cetyltrimethylammonium bromide (CTAB) DNA extraction protocol. For cell lysis, 5ml CTAB buffer (0.1 M Tris, 0.7 M NaCl, 10 mM EDTA, 1% CTAB, 1% beta-mercaptoethanol) was added into 15 ml tube containing sample. After incubation at 65°C for 30 min, an equal volume of chloroform was added. The mixture was centrifuged at 10,000 rpm for 10 minutes and the supernatant was mixed with an equal volume of isopropanol. After centrifugation at 10,000 rpm for 30 minutes at 4°C, the supernatant was discarded and the pellet was washed twice with 70% and 90% ethanol. DNA was eluted with 50 µl elution buffer (Qiagen).

Internal transcribed spacer (ITS) and 16S rRNA amplicons were generated using barcode primer pairs ITS3ngs(mix)/ITS4^73^ and V3/V4^74^, respectively. Amplicon levels were standardised using the SequalPrep Normalization Plate 96 Kit (Invitrogen Corporation, Carlsbad, CA, USA, Cat. #A10510-01). Concentration of pooled and standardised amplicons was performed using Agencourt AMPure XP beads (Beckman Coulter, Brea, CA, USA, Cat. #A63881). All amplicon libraries were sequenced with Illumina MiSeq PE300 using 2 x 300 bp paired-end chemistry performed by the NGS High Throughput Genomics Core at the Biodiversity Research Center, Academia Sinica, Taiwan.

### Construction of OTU table from amplicon reads

Samples were demultiplexed using sabre with one nucleotide mismatch (v1.0; https://github.com/najoshi/sabre) with their respective barcodes. Adaptor and primer sequences were trimmed using USEARCH^75^ (v10). Sequence reads were processed according to the UPARSE^76^ pipeline. Forward and reverse reads were merged and filtered using USEARCH. Operational taxonomic units (OTUs) were clustered at 97% sequence identity with a minimum of 8 reads per cluster to denoise the data and remove singletons. The OTU table was generated using the *usearch_global* option and analysed with the *phyloseq*^77^ package (v1.28.0) detailed in **Supplementary Information**. SINTAX^78^ algorithm was used to classify the OTU sequence taxonomy against the RDP^79^ training set (v16) and the UNITE^80^ database (v7.1).

### Morphology observation of *D. spatulata* stalk gland

The stalk glands of *D. spatulata* were photographed using an Olympus microscope (Olympus CX31) and digital camera (Nikon D7000). Staining was performed using cotton blue reagent as described^81^. Scanning electron microscopy (SEM) of the *D. spatulata* stalks were prepared as follows. Leaves tissues were fixed at 4 degrees for 1 hour with P4G5 solution (4% paraformaldehyde and 2.5% glutaraldehyde in 0.1 M phosphate buffer). After three washes with 0.1 M phosphate buffer, secondary fixation was performed in a 1% solution of osmium tetroxide for 2 hours at room temperature. The fixed samples were passed through a series of dehydration step in 30, 50, 70, 80, 90, 95, 100, 100 and 100% alcohol before drying in a critical point dryer (Hitachi model HCP-2). The dried samples were then coated with a layer of gold using a sputter coater (Cressing-ton model 108). SEM observations of *Drosera* leaves grown under different conditions (laboratory condition, inoculated with *A. crateriforme* and from wild) were performed on a JSM-7401F scanning electron microscope (JEOL) at the Institute of the Plant and Microbial Biology, Academia Sinica, Taiwan.

### Inoculation experiment

We used *D. spatulata* plants derived from tissue culture in vermiculite with ddH_2_O as our control samples. We inoculated *A. crateriforme* and *Ph. herbarum* on *D. spatulata* leaves and incubated the plants at 25 degrees for 30 days. Photographs of *D. spatulata* in different treatments including control plants were taken daily and the area of leaf displaying symptoms of infection was quantified every two days using ImageJ^82^. Whole *D. spatulata* plants were dried with tissue paper and weighted on the first and 30^th^ day of the experiment.

### Feeding experiment of *D. spatulata*

10^-5^ g of sterilised ant powder was added on a single leaf of an individual *D. spatulata* plant with or without *A. crateriforme* inoculated. The capture time (from the time ant substrate was added until the tentacles fully covered the prey) and digestion time (from the time tentacles fully covered the prey until the trap reopened) of five replicates were recorded. The same set of experiments were repeated with additional application of Protease Inhibitor Cocktail (#P9599, Sigma). To compare the effect of protein digestion in mucilage from *D. spatulata* with and without *Acrodontium*, mucilage from 30 plants in each treatment were collected prior to the experiment. We mixed mucilage with BSA labelled with HSP-biotin at both 25 degrees for 16 hours and 24 hours. Western blot was made (**Supplementary Information**) and the amount of protein digestion was quantified using ImageJ^82^.

### Comparative genomics and phylogenomics

The assembly and annotation of *A. crateriforme* are detailed in the **Supplementary Info**. A total on 32 genomes from representative fungi and plants were downloaded from JGI and NCBI databases (**Supplementary Table 5**). For each gene, only the longest isoforms were selected for subsequent analysis. Orthogroups (OGs) were identified using Orthofinder^83^ (ver. 2.5.5). For each orthogroup, an alignment of the amino acid sequences each gene was produced using mafft^84^ (version 7.741). A maximum likelihood orthogroup tree was made from the alignment using IQtree^85^ (version 2.2.2.6). A species phylogeny was constructed from all orthogroup trees with ASTRAL-III^86^ (ver. 5.7.1). OG gains and losses at each node of the species phylogeny were inferred using DOLLOP^87^ (ver. 3.69.650).

### Transcriptome analysis

Total RNA of were extracted from *D. spatulata* and underwent transcriptome sequencing (**Supplementary Info**). RNA-Seq raw reads were trimmed using fastp^88^ (v0.23.2) to remove the adaptor and low-quality sequences. The trimmed reads were mapped to the corresponding genome using STAR^89^ (v 2.7.10b) and assigned to gene count using featureCounts^90^ (v 2.0.3,). Notably, the reads from coexistence treatment, with or without ant powder, were mapped to both *A. crateriforme* and *D. spatulata* genome^18^. To prevent the false positive of gene expressions, sequences mapped to both genomes and had low mapping qualities were excluded from further analyses. The samples of *D. spatulata* exposed to ant powder which obtained in two different time points were grouped as digestion sample. The differentially expressed genes (DEGs) of different conditions comparing to control, were inferred by DESeq2^91^ (v1.38.3; padj < 0.05 & |log2FD| > 1). The gene ontology enrichment of comparisons was identified using topGO^92^ (v2.50.0). We also performed weighted gene co-expression network analysis (WGCNA) to further categorise the expression patterns of peptidases respectively in *A. crateriforme* and *D. spatulata*. Due to the present of peptidase without any expressions across conditions in *D. spatulata*, we removed the 30% lowest-expressed genes in each transcriptome using the sum of samples. The descriptions and annotations of every DEG in *A. crateriforme* and *D. spatulata* are available in **Supplementary Table 10 and 11**, respectively.

### Phytohormone analysis

We loaded different treatments (1g/L of BSA, chitin, BSA+chitin, and ants, 10^6^ spores/ml of *A. crateriforme*, and 10^5^ spores/ml of *A. crateriforme*) to *D. spatulata* leaves for 2 hours. Then, leaves were cut and washed in de-ionized water to remove residue. Then, leaves were rapidly frozen using liquid nitrogen. The time from cutting to freezing consistently remained under 30 seconds. The prepared samples were then ready for the metabolite extraction.

The metabolite extraction used 1 mL CHCl_3_:MeOH (2:1) as extraction solvent with Dihydrojasmonic acid (H_2_JA) (7.5 ng for 0.3 g leaf tissue) adding as internal standards. Equal volumes of the supernatant were stored at −80 °C. Samples were reconstituted in 50 µL of 20 % aqueous methanol each. The samples were analysed by the Vanquish UHPLC system coupled with a Dual-Pressure Ion Trap Mass Spectrometer (Velos Pro, Thermo Fisher Scientific). Jasmonoyl-L-isoleucine (JA-Ile) and its standard H_2_JA were separated by an HSS T3 column (Waters ACQUITY HSS T3 100Å, 1.8 µm, 100 × 2.1 mm) at 40°C using the mobile buffer consisted of 2% ACN/0.1% FA (Buffer A) with an eluting buffer of 100% ACN/0.1% FA (Buffer B) with a 11 min gradient of 0.5-30% Buffer B at 0-6 min, 30-50% B at 6-7 min, 50-99.5% B at 7-7.5 min, 99.5-0.1% B at 9.5-10 min and then equilibrated by 0.1% B at 10-11 min. The selected m/z 322.20 to 130.09 for JA-Ile and 211.13 to 59.01 for H_2_JA^93^.

### Data Availability

All sequences generated from this study were deposited on NCBI under BioProject PRJNA1034788 and accession numbers of individual samples can be found in **Supplementary Table 1 and 12**.

## Funding

This work was supported by National Science and Technology Council (112-2628-B-001-005-) to IJT. PFS is supported by the doctorate fellowship of the Taiwan International Graduate Program, Academia Sinica of Taiwan.

## Acknowledgements

We thank Ting, See-Yeun, Shen-Feng Sheng, Chia-Lin Chung for initial discussions and encouragements. We thank Ko-Hsuan Chen for reading and commenting of the manuscript. We thank Hsin-Han Lee for producing an initial version of the *A. crateriforme* assembly. We thank Mitsuyasu Hasebe for advice on experiments on *Drosera spatulata*.

## Author contributions

IJT conceived the study; PFS carried out the sampling and the experiments with guidance from YFL, HMK, YCJL and YLC. PFS performed and analysed the amplicons with guidance from YFL and DZH. MJL sequenced the amplicons. PFS and RK identified the *A. crateriforme* strains. HMK carried out ONT sequencing. IJT carried out the *A. crateriforme* genome assembly and annotation. PFS, MRL, YCL, and IJT carried out comparative genomic and transcriptomic analyses. PFS, IFW and YLC quantified the phytohormones. PFS and IJT wrote the manuscript with input from MRL, YCL, RK, YCJL, YLC and others. All authors read and approved the final manuscript.

## Notes

### Competing Interest Statement

The authors have declared no competing interest.

