## Supplementary Info for "An acidophilic fungus is integral to prey digestion in a carnivorous plant"

**Extended Data Fig. 1 –** Scanning electron microscope (SEM) image of sundew leaves **a-c.** under sterilised conditions, **d-f.** inoculated with *A. crateriforme*, and **g-i.** collected from wild.

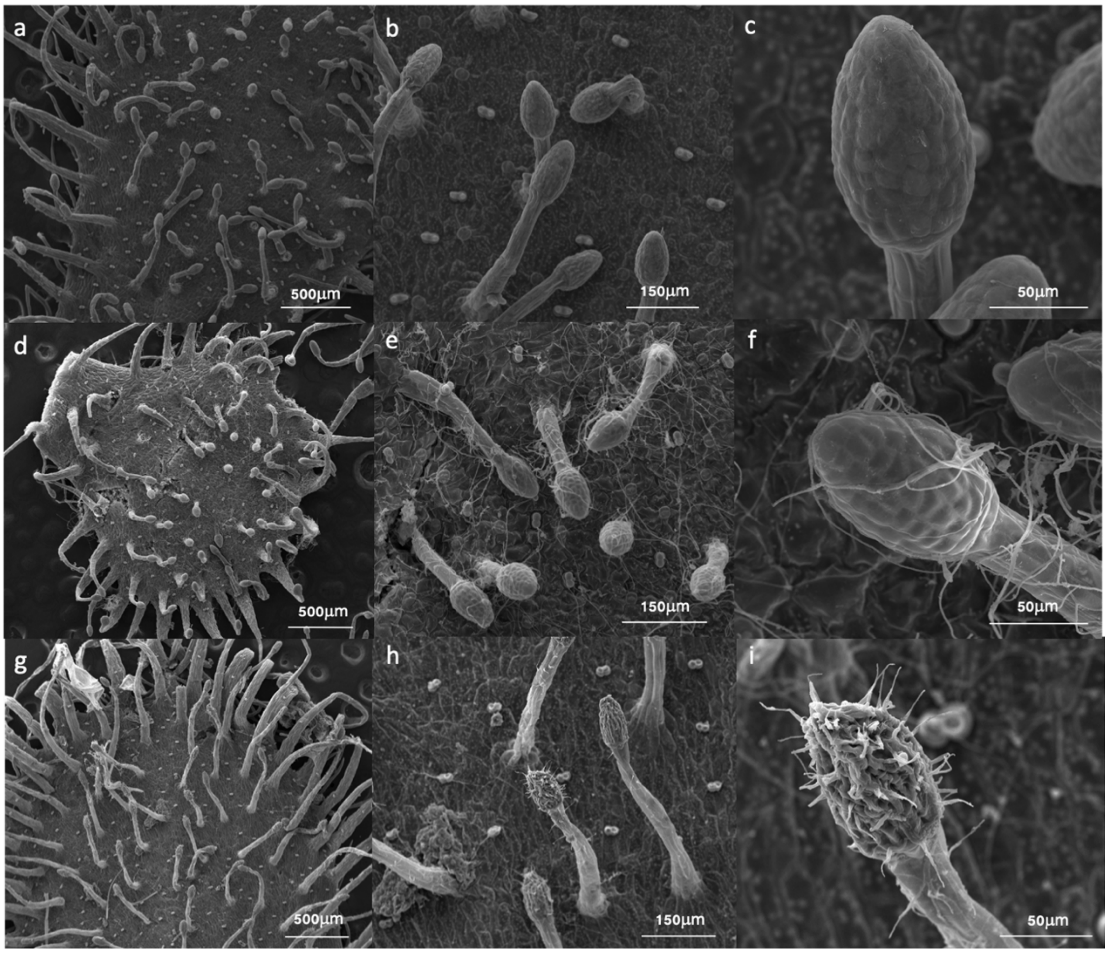

**Extended Data Fig. 2 –** Fungal growth on remains of dead arthropod (covered by evenly spread hairs) on wild *Drosera spatulata*. Fungal hyphae indicated by white arrowheads, two conidiophores of *Acrodontium crateriforme* by red arrows.

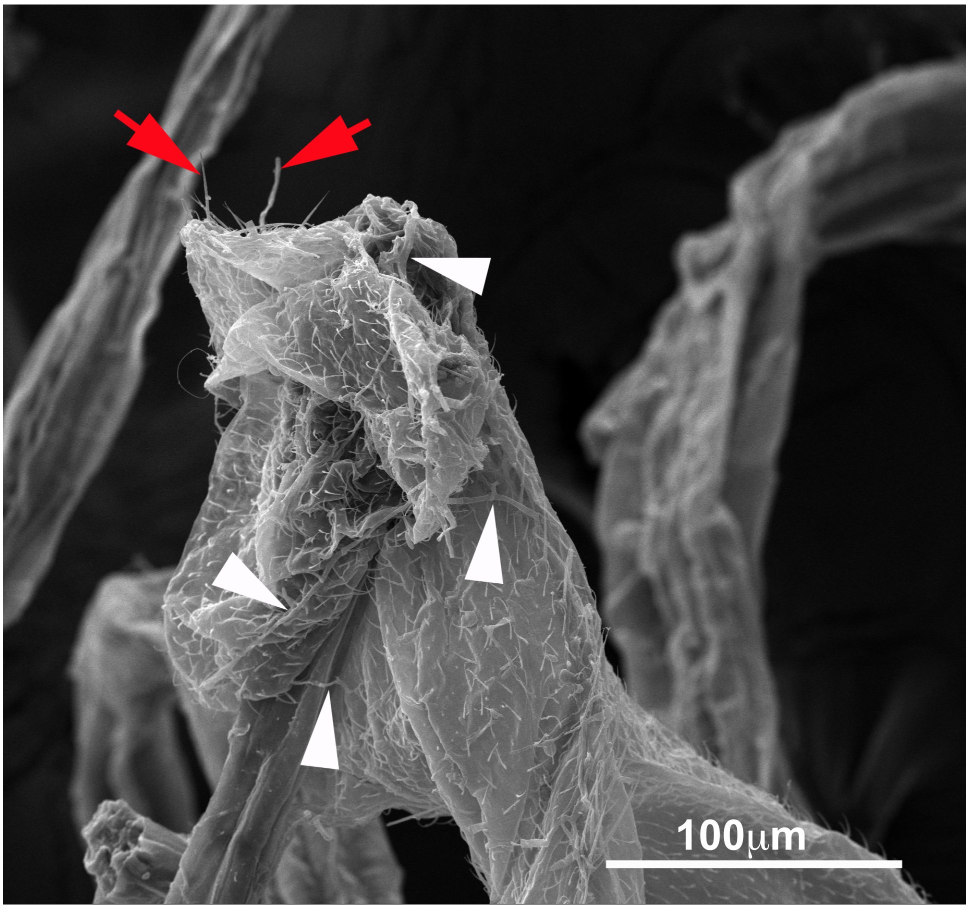

### Supplementary Methods

#### Collection of sundew mucilage and surrounding plants

In 2018-2019, we collected *D. spatulata* mucilage samples from five collection sites located in Northern Taiwan (**Supplementary Table 1**). Each sample is a pooled mucilage from 30 *D. spatulata* leaves of the same site by using filter paper of size 1 cm x 1.5 cm. The fresh leaves of plants and mosses surrounding *D. spatulata* were also wiped and considered as environmental samples. To determine the temporal microbial dynamics in mucilage, we further sampled Shumei and Shuangxi sundew mucilage monthly from June 2017 to April 2018 except two time points. To understand the spatial extent of plant-fungus coexistence, we collected 52 mucilage samples from 17 additional sites.

#### Amplicon analysis

Prior to data analysis, the accuracy of the UPARSE pipeline was assessed using positive controls containing known species. Sequence samples were decontaminated by R package *decontam*[1] and analysed with the *phyloseq*[2] package (v1.28.0). Unclassified ITS and 16S operational taxonomic units (OTU) sequences were identified by BLAST against the NCBI *nr* database. OTUs of non-fungal and non-bacterial origin were removed from subsequent analyses. Differentially abundant taxa were determined using DESeq2[3] (v1.24). Amplicons were reported at the genus level. A total of 15,366,944 fungal and 3,623,798 bacterial sequence reads were assigned to 5,760 fungal and 3,085 bacterial OTUs, respectively. After rarefaction, each fungal and bacterial sample had 14,000 and 15,000 sequence reads, respectively. One sample without this sequence depth was removed.

#### Isolation and identification of fungal species from sundew mucilage

Mucilage-soaked filter papers were placed in potato dextrose broth (PDB) containing chloramphenicol (50ug/ml) and incubated for at 30°C for 24 hours. The broth culture was then serially diluted ($10^{-2}$, $10^{-3}$, $10^{-4}$) and 200 μl of the diluted culture was transferred to potato dextrose agar (PDA). The plate was spread on sterilised glass until the medium was dry and incubated at 30°C for 24 hours. Single colonies were transferred onto fresh PDA using sterilised toothpicks and examined for morphology using light microscopy. Genomic DNA was extracted from the fungal pure cultures corresponding to the morphological description of *Acrodontium*, followed by amplification and sequencing of the fungal ITS region as previously described[4,5]. To construct a phylogenetic tree, ITS sequences of the sequenced 18 isolates, 15 *Acrodontium, Teratosphaeria biformis* and *Aureobasidium pullulans* ITS sequences from NCBI were first aligned using MAFFT[6] (v.7.4) and trimmed using trimAl[7] (v.1.2). A maximum likelihood phylogenetic tree with 100 bootstraps was produced from the alignment using IQtree[8] (v.1.6.1). All 18 isolates were classified as *A. crateriforme* based on morphological description and grouped with the *A. crateriforme* ITS sequence.

**Preparation of sterilised *Drosera spatulata***

*D. spatulata* seeds were collected from the sampling sites and sterilised by washing with 70% ethanol for 10 seconds, 3% (w/v) calcium hypochlorite ($CaCl_2O_2$) for 30 seconds and finally rinsed 3 times with dd$H_2O$. Surface sterilised seeds were pregerminated on 0.5% water agar and incubated in the dark at 20°C for 6-8 weeks. Shoots were then transferred to 1/2 MS agar with pH adjusted to 5.7. Shoots were grown under white fluorescent light at 20°C with a 16h:8h, light:dark photoperiod for 90 days as recommended[9].

***A. crateriforme* growth conditions in different condition**

*A. crateriforme* was inoculated in nutrient-rich medium agar plate in different temperatures (20, 25, 30 degree) and pH value (pH 3-pH 9) for two weeks. For testing *A. crateriforme* growth conditions in different ants, we supplemented ant powder in nutrient-rich or nutrient-poor medium agar plate. Then, *A. crateriforme* was inoculated in these medium with or without ant powder for two weeks. The fungal growth area was photographed using Nikon digital camera every two days. Then, we measure growth area by using ImageJ [10].

**Genome sequencing, assembly and annotation of *A. crateriforme***

Genomic DNA was subjected to Oxford Nanopore library preparation according to the manufacturer's instructions (SQK-LSK109), and sequenced on a GridION instrument. Basecalling was using Guppy (ver. 3.0.3). For Illumina sequencing, genomic DNA was used for NEB Next Ultra library preparation and 150bp paired-end reads were generated on a Novaseq 6000 sequencer. The nanopore reads was first corrected from the initial assembly of the canu[11] assembler (ver. 1.9), which were then assembled using the flye assembler (ver. 2.5). The initial assembly was polished by Racon[12] (four iterations; ver. 1.4.11), followed by Medaka (ver. 0.11.0; https://github.com/nanoporetech/medaka) using nanopore reads and Pilon[13] with Illumina reads. The mitochondrial genome was assembled separately using NOVOPlasty[14] (ver. NOVOPlasty2.7.0.pl).

**Gene prediction of *A. crateriforme***

The transcriptome reads were mapped to the *A. crateriforme* genome assembly using STAR[15] (ver. 2.7.7a) and assembled using Trinity[16] (ver 2.13.2; guided approach), Stringtie[17] (ver 2.1.7) and Cufflinks[18] (ver 2.2.1). Transcripts generated by Trinity were mapped to the assembly using Minimap2[19] (ver 2.1, options: -ax splice), and splice junctions were quantified using Portcullis[19] (ver 1.2.3). The gene predictor Augustus (ver 3.4.0) and gmhmm[20] were trained using BRAKER2[15,21] (ver. 2.1.6) and SNAP[22] with proteomes and RNAseq mappings as evidence hints to generate an initial set of annotations. The assembled transcripts selected by MIKADO[23] (ver 2.3.3), proteome downloaded from Uniprot Fungi (version October 2019) as homology, and BRAKER2 annotations were combined as evidence hints for input into the MAKER2 annotation pipeline[24] to produce a final annotation for each species. Repetitive elements were identified based on the protocol by Berriman *et al*. [25], and masked using Repeatmasker[26] (ver 4.1.2).

Functional domains within the proteomes were identified using pfam_scan[27] (ver. 1.6) against the downloaded Pfam database[28] (ver. 36). Diamond[29] (ver. 2.1.6) was utilised to identify transporters, blasting proteomes against TransportDB[30] (ver. 2.0). Proteomes were functionally annotated to identify carbohydrate-active enzymes (CAZy) and peptidases using dbCAN[31] (ver. 2.0.11), and MEROPS[32] (ver. 12.4), respectively. Annotations of biosynthetic gene cluster (BGC) regions and Gene Ontology (GO) terms were performed using antiSMASH[33] (fungi ver. 7.0.1) and eggNOG[34] (ver. 2.1.12), respectively.

Analysis and visualisations were conducted under R[35] environment (ver. 4.3.1). Various packages were utilised to enhance the analysis: topGO[36] (ver. 2.52.0) for GO enrichment analysis, pheatmap[37] (ver. 1.0.12) and ggplot2[38] (ver. 3.4.4) for gene expression visualisation.

**Greenhouse experiment for plant-microbe interaction**

Six treatments (tissue culture of *D. spatulata* w/o predation, *A. crateriforme* only w/o predation and co-culture of *D. spatulata* and *A. crateriforme* w/o predation) are used for plant-fungal interaction analysis. For the tissue culture samples, we transfer the tissue-cultured *D. spatulata* from the 1/2 MS medium to vermiculite with ddH2O and incubate for 3 weeks. For fungal inoculation, removed the conidia from 14-day fungal colonies by washing sterile distilled water. Then, adjusted the suspension to $10^6$ spores/ml. After that, use pipette to inoculate microbe into stalk gland. After 2 weeks following inoculation, sundew is able for the experiment. We added insect substrate to five leaves on an individual plant and harvested them as predation samples after 72 hours.

**RNA extraction**

We pooled 80 leaves as one replicate and each treatment had five replicates. At each harvest, leaf tissue was cut from the plant and washed immediately in de-ionized water to remove prey residue. The leaf was then flash-frozen in liquid nitrogen (−196 °C). The time from cutting to freezing was always <30 s. Plant and fungal RNA is extracted by modified CTAB method. 2ml CTAB buffer (0.1 M Tris, 2 M NaCl, 25 mM EDTA, 2% CTAB, 1% PVP-40, 2% beta-mercaptoethanol) was added into 2 ml tube containing leaves sample. Samples were frozen in liquid nitrogen and grinding by Precellys 24 tissue homogenizer. After incubation at 65°C for 20 min, an equal volume of chloroform: isomylalcohol (24:1) was added. The mixture was centrifuged at 12,000 rpm for 10 minutes twice. Supernatant were mixed 1/3 volume of LiCl and put in 4 °C overnight for RNA precipitate. After centrifugation at 10,000 rpm for 30 minutes at 4°C, the supernatant was discarded and the pellet was washed twice with 70%. RNA was eluted with 30 µl DEPC water. RNA samples were sequenced by using Novaseq 6000.

**Western blot**

We mixed 10 µl protein sample with 10 µl of 2X loading dye in a 0.2 µl PCR tube. The samples are then incubated for 10 minutes at 99 degrees in the PCR machine. After incubation, protein samples were loaded onto the SDS-PAGE. Electrophoresis used 90 V for 20 minutes in the stacking gel and then used 120 V for 120 minutes in the running gel. After electrophoresis, we transfer the SDS-PAGE into a 1X transfer buffer for 10 minutes. Subsequently, we put soaked filter papers, membrane, and SDS-PAGE together. The machine parameters for transferring were set to 0.8 mA/cm² for 1 hour. We washed the membrane in 1x PBST for 5 minutes twice. Then, soaked the membrane in blocking buffer at room temperature for 1 hour. Then, washed membrane in 1x PBST for 5 minutes twice. To proceed, we add the secondary antibody solution and shake it for 30 minutes at room temperature. After that, we washed the membrane with 1X PBST for 8 minutes at room temperature three times. Finally, we prepare the Luminol substrate and peroxidase in a 1:1 ratio (750 μl + 750 μl). The membrane is placed into the detecting chamber for signal detection and the detection is performed by the machine.

### Supplementary Figures

**Supplementary Fig. 1. *Drosera* spatulata in its natural habitat. a.** *D. spatulata* typically grows on the cliffs. **b.** When zoomed in, *D. spatulata* is surrounded by grasses and moss.

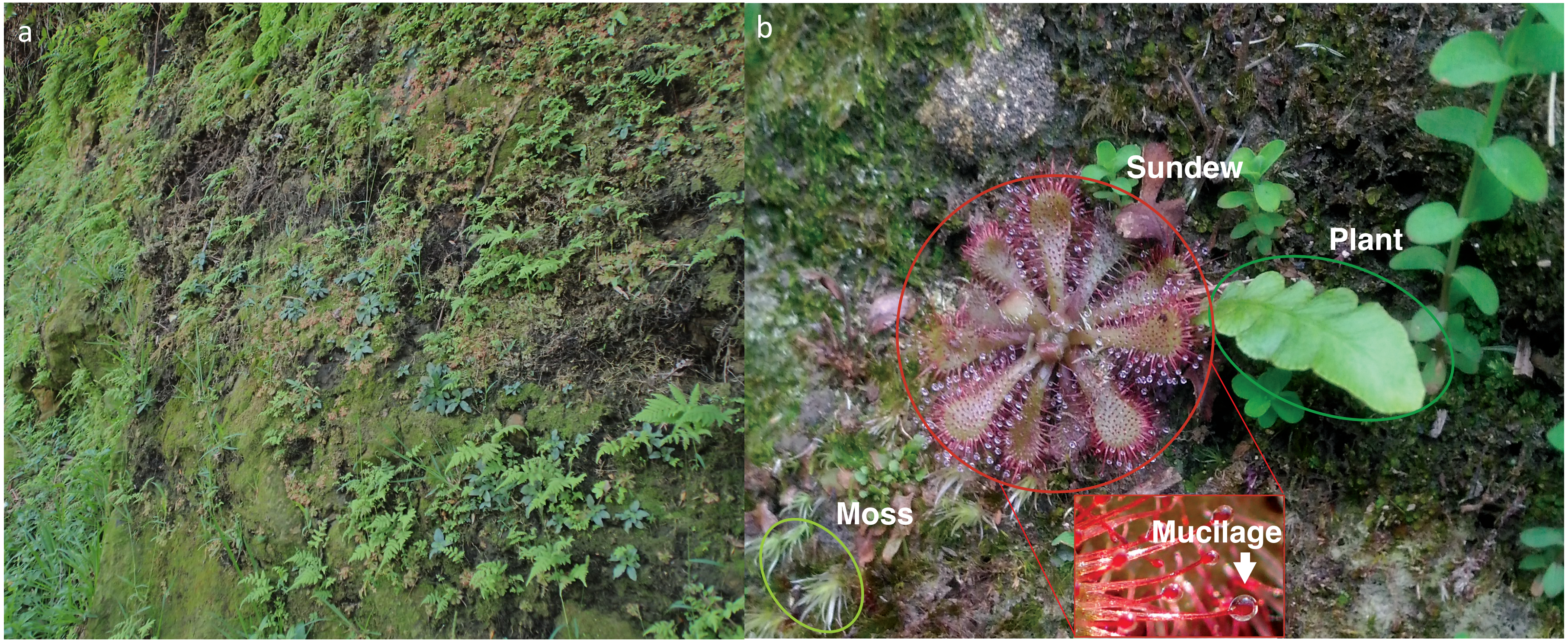

**Supplementary Fig. 2. Beta diversity (Bray-Curtis index) of fungal communities after the dominant *Acrodontium* OTU was excluded.** Ellipses show the 95% confidence intervals for the mean of samples of the same source and site.

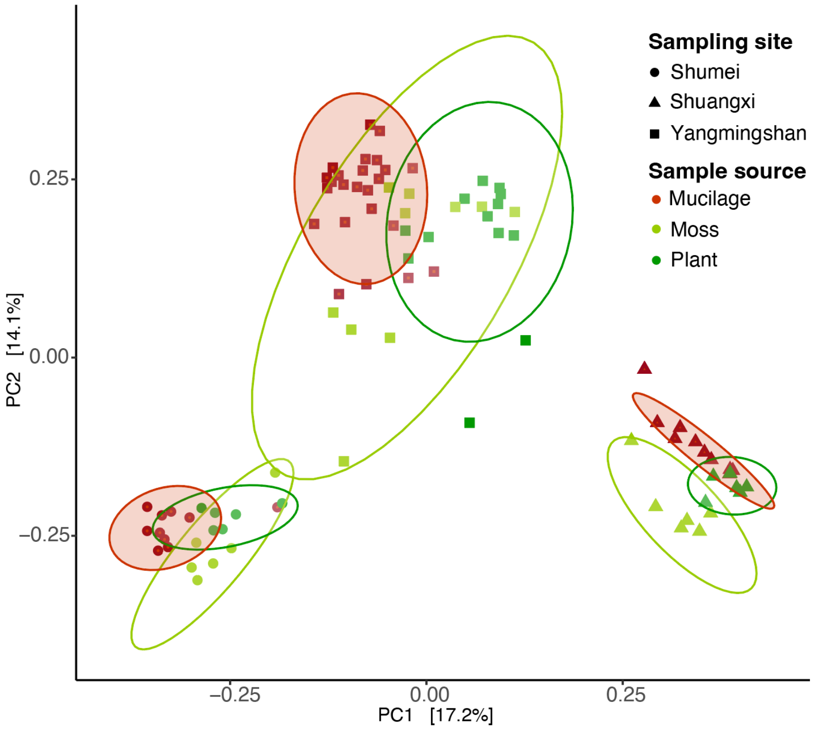

**Supplementary Fig. 3. Distribution of *Acrodontium* genus from Globalfungi.**

**a.** Globalfungi showing *Acrodontium* mostly located in the forest followed by grassland habitat. **b.** Relative abundance of *Acrodontium* in different location.

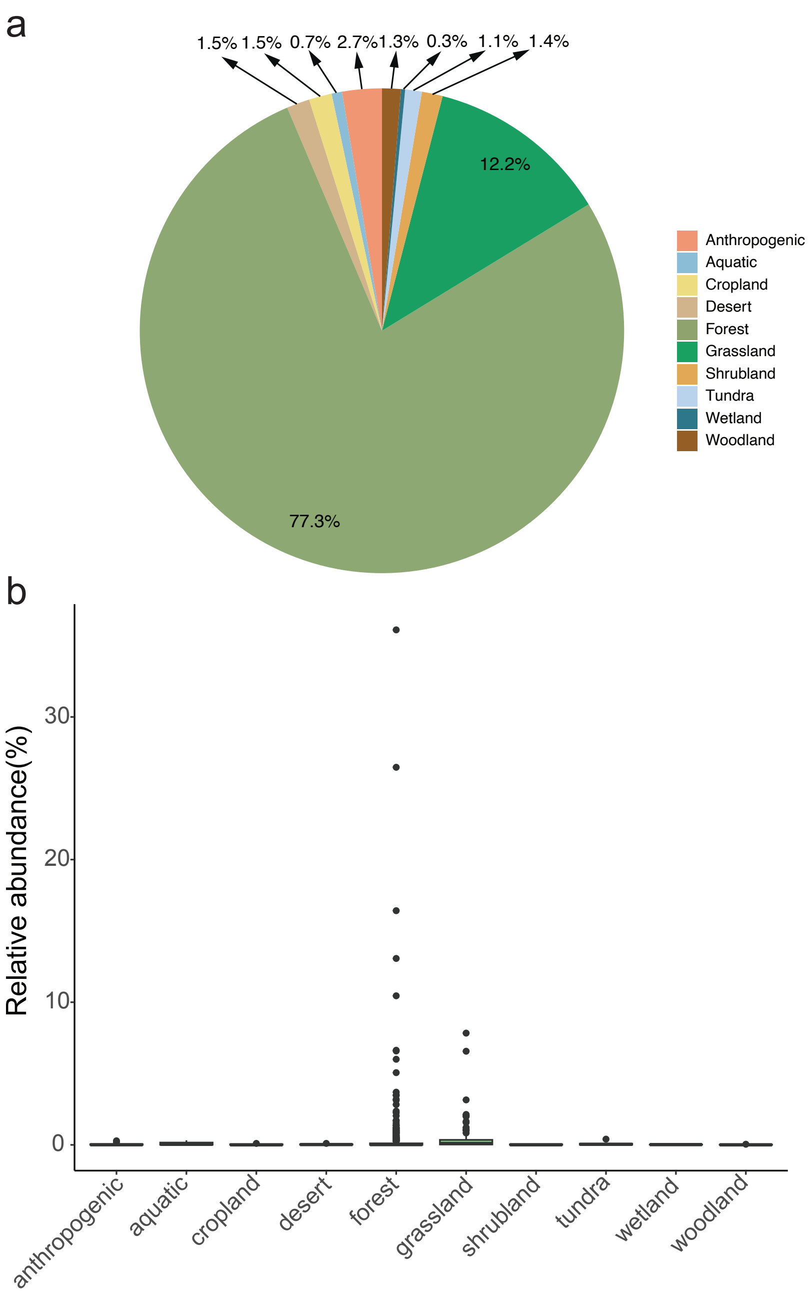

**Supplementary Fig. 4 – *Acrodontium* genus preferred pH value from Globalfungi data. a.** Most samples harbouring the *Acrodontium* OTU did not have information on pH. Based on available information, *Acrodontium* genus was preferentially isolated in samples of pH 4-5. **b.** Relative abundance of *Acrodontium* in samples of different pH.

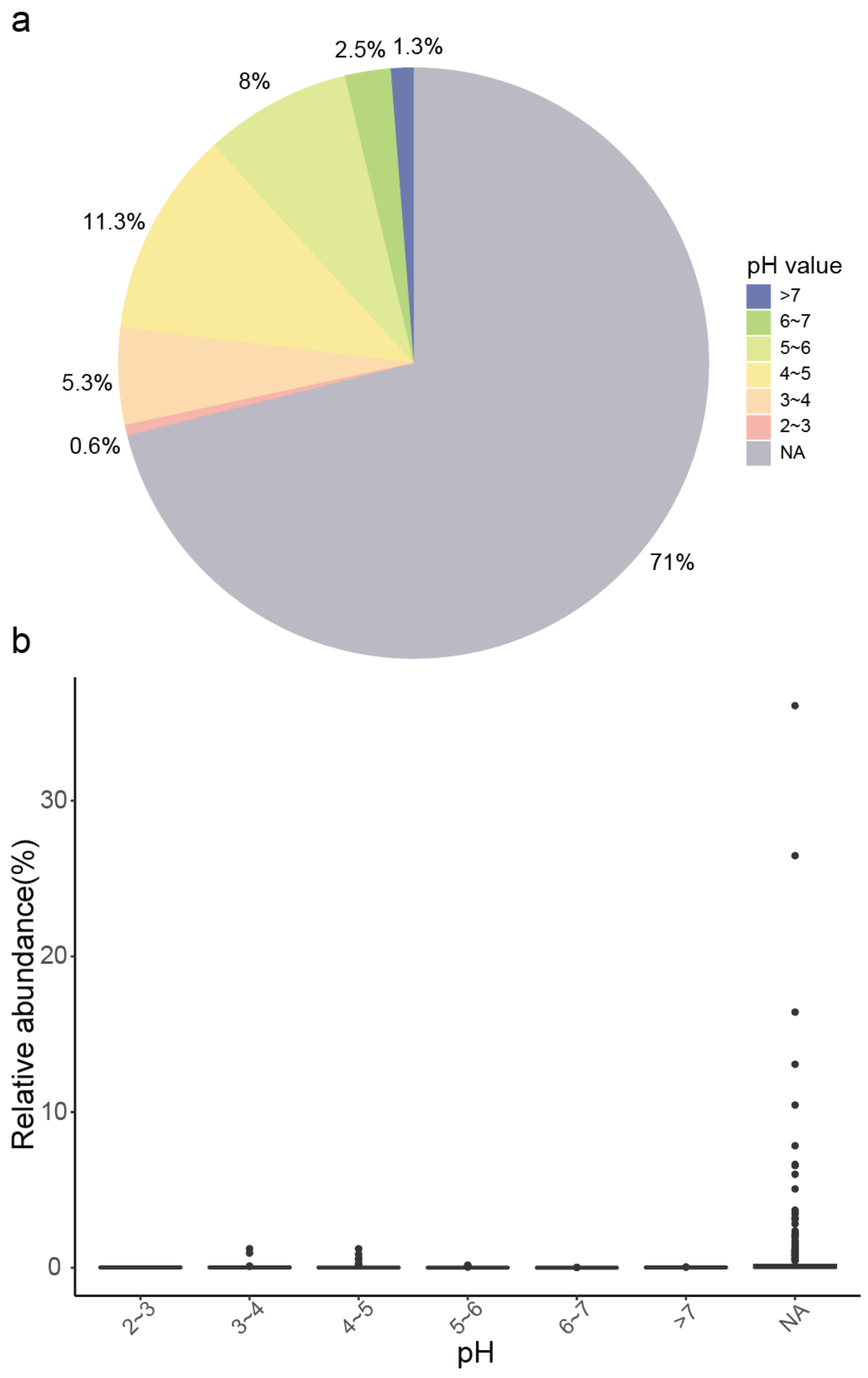

**Supplementary Fig. 5. ITS phylogeny of *Acrodontium crateriforme.* a.** Photo shows the morphology of *Acrodontium crateriforme* grown in Potato dextrose agar. **b.** Red colour represents the consensus sequence of the *Acrodontium* OTU from amplicon data. Brown colour represent sequences of *A. crateriforme* strains that were isolated from *Drosera spatulata* mucilage

a b

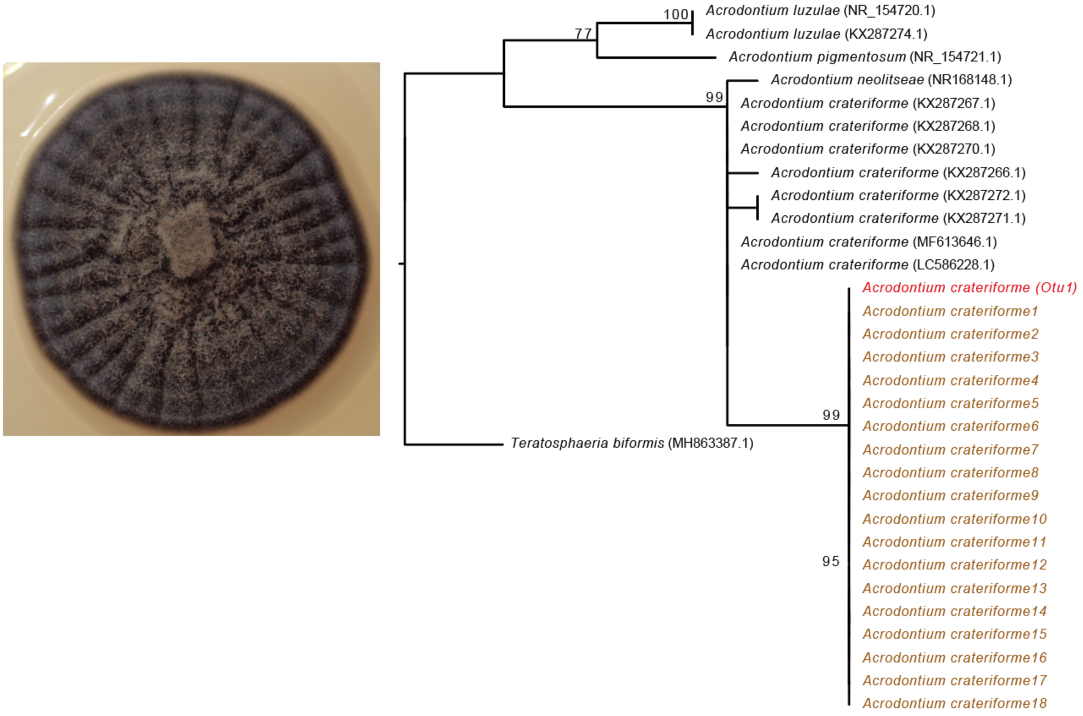

**Supplementary Fig. 6**. **ITS phylogeny of *Phoma herbarum.*** Photo shows the morphology of *Phoma herbarum* grown in Potato dextrose agar. Red color represents *Phoma herbarum* from amplicon data. Brown colour represent *Phoma herbarum* we isolated from *Drosera spatulata* mucilage

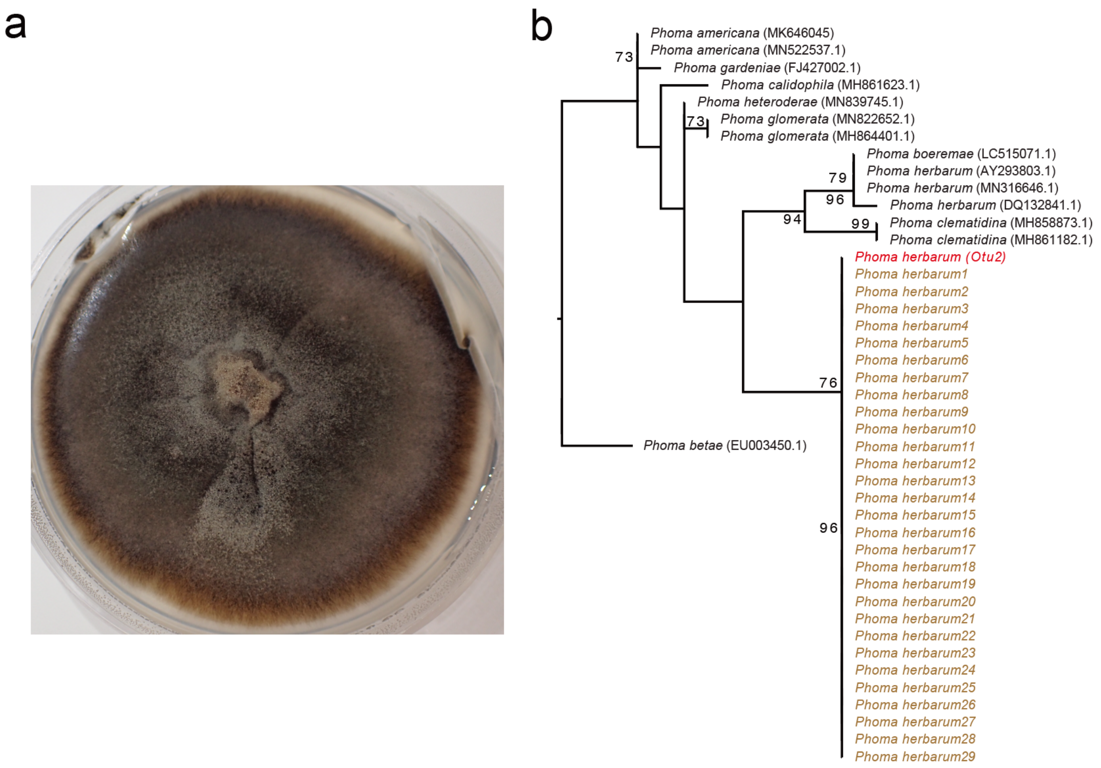

**Supplementary Fig. 7. The growth profile of *A. crateriforme* and *P. herbarum* in different pH value.**

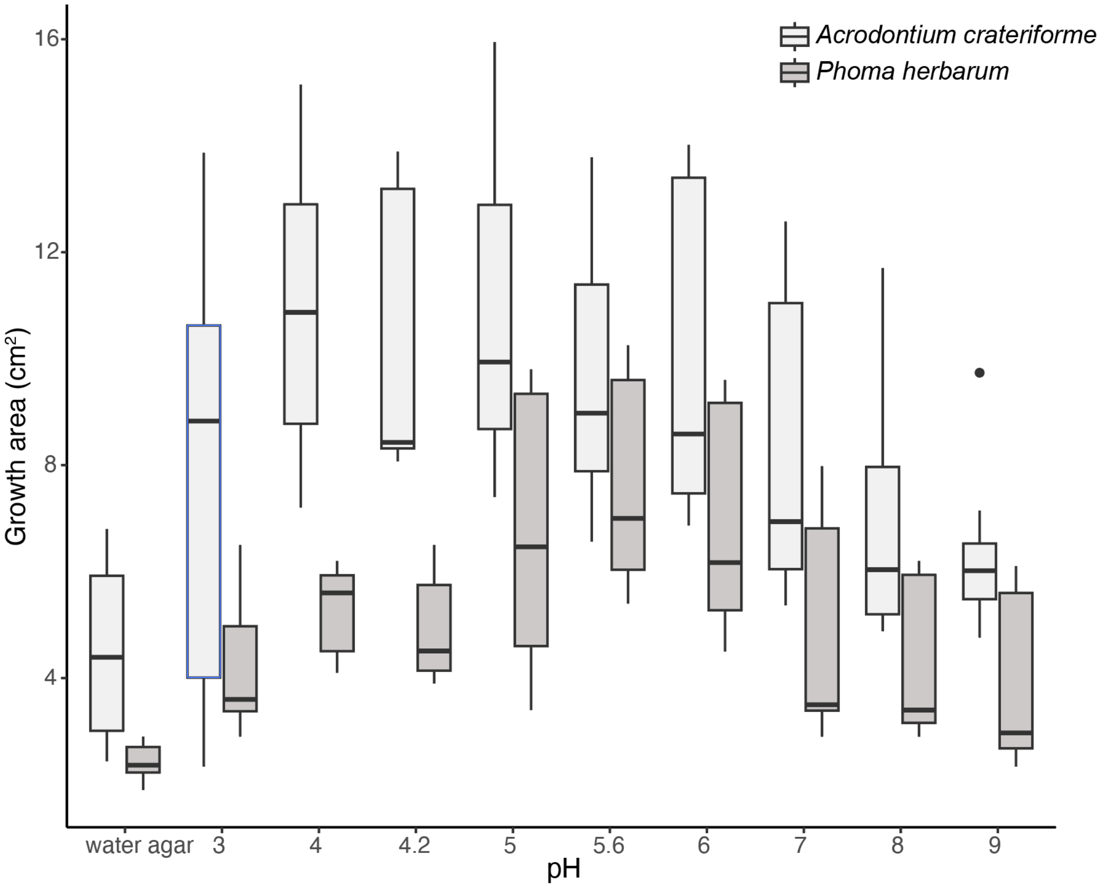

**Supplementary Fig. 8. The growth profile of *A. crateriforme* and *P. herbarum* in different temperatures.**

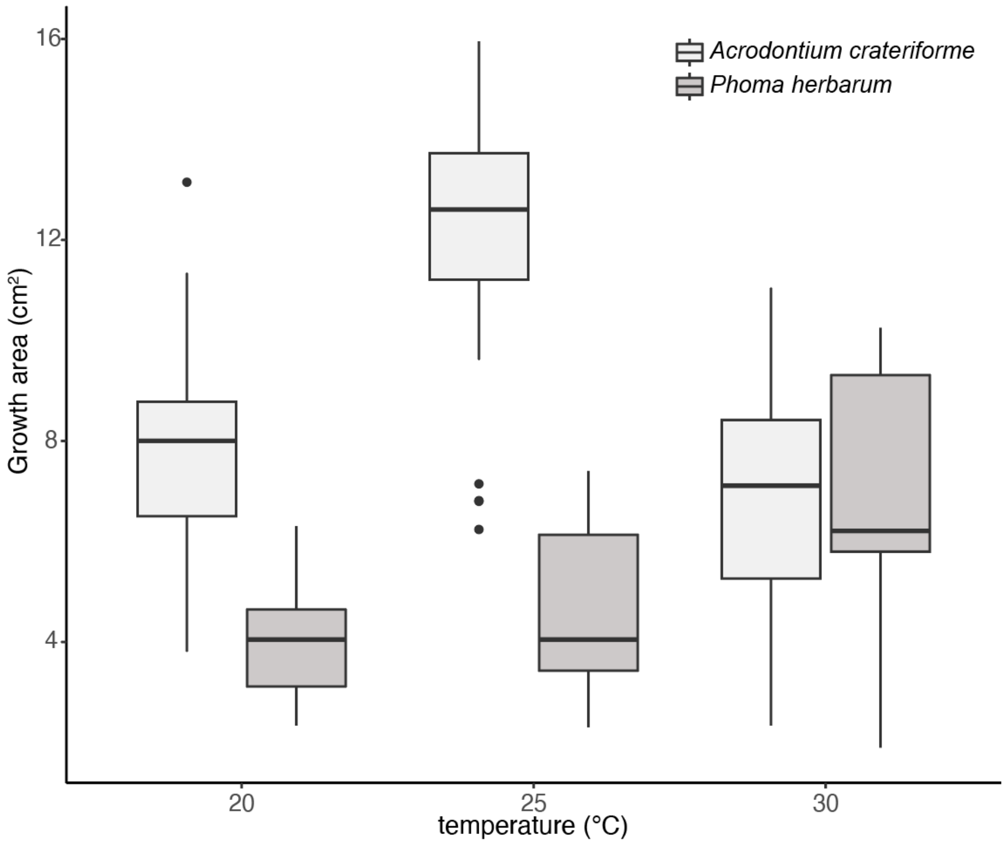

**Supplementary Fig. 9 Temperature data in Shumei and Shuangxi from Taiwan Central Weather Administration.**

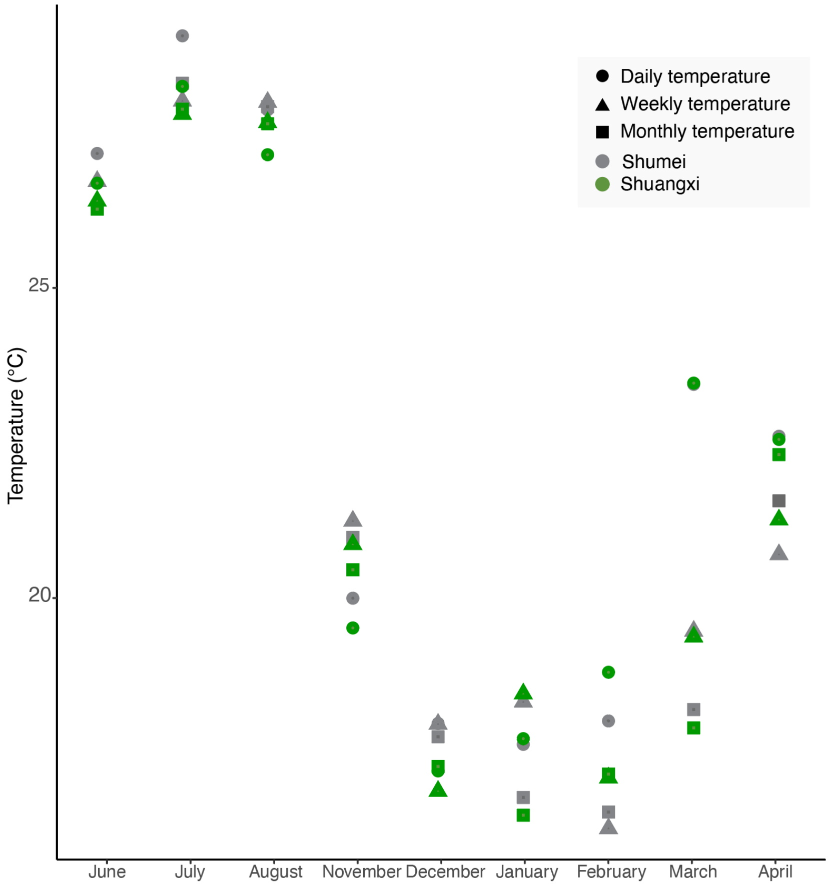

**Supplementary Fig. 10 Growth profiling of *A. crateriforme* in different media condition**

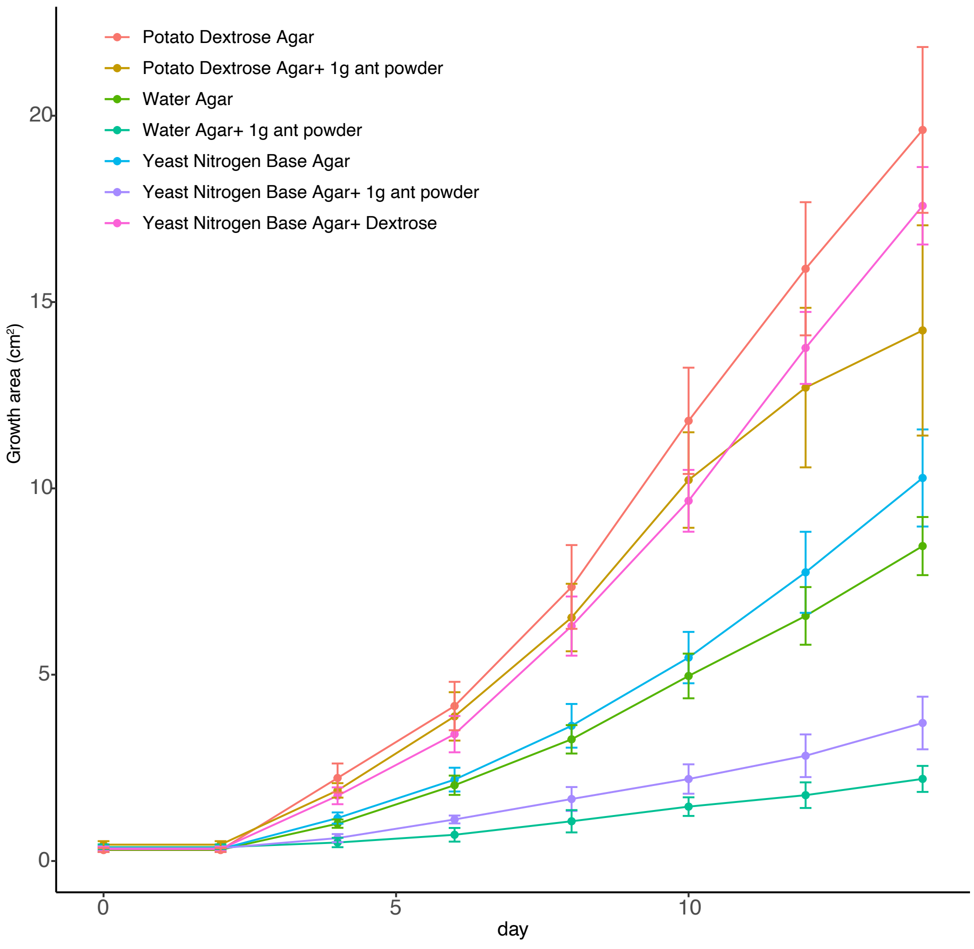

**Supplementary Fig. 11 Weight of different fungi inoculated in *Drosera spatulata.*** Boxplots show *Drosera spatulata* weight. Different colors show the weight at day 0 and different treatment of inoculation after a month (30 days).

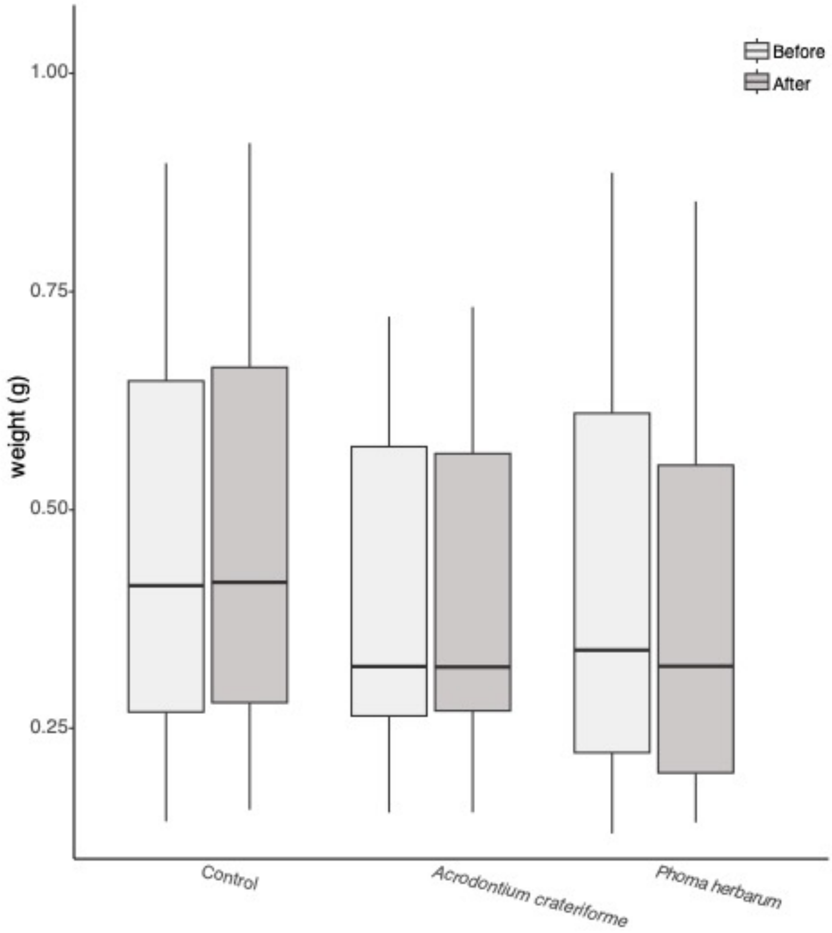

**Supplementary Fig. 12 Western plot of biotin-BSA digestion in mucilage under different treatment.**

Unstained protein ladder: Thermo 26630, Prestained protein ladder: Thermo 26616. Negative control indicated pH4 MES buffer. Positive control indicated biotin-BSA in pH7 or pH4 of MES buffer. *Drosera* mucilage + biotin-BSA shows *Drosera* mucilage mixed with biotin-BSA in pH4 MES buffer. *Drosera* mucilage shows *Drosera* mucilage in pH4 of MES buffer

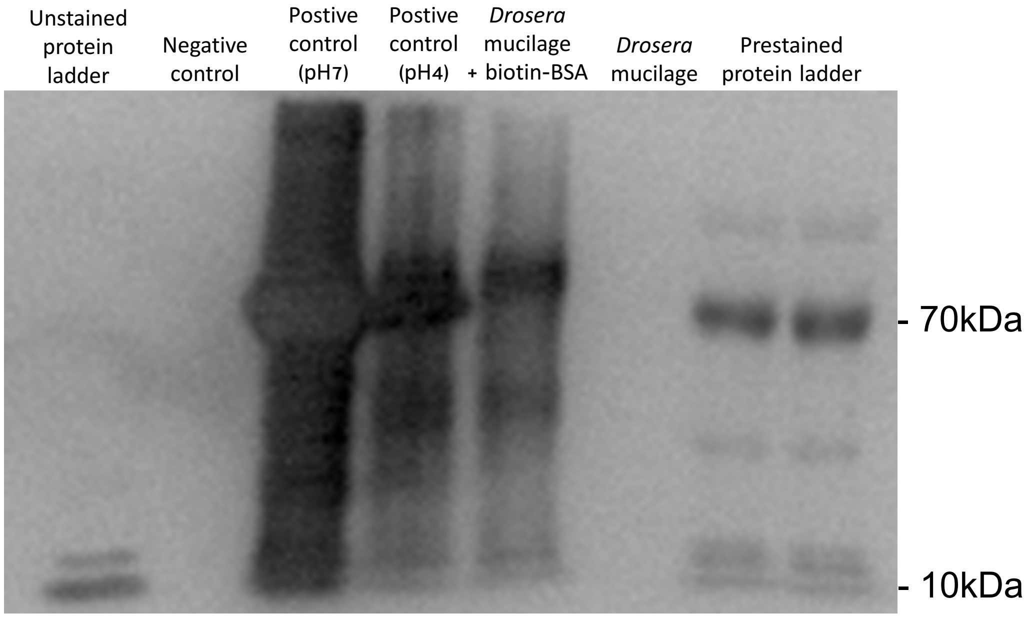

**Supplementary Fig. 13**. **PCA of protein family domain numbers from 25 fungal species a.** PCA of all species, **b.** zoomed in plot. Species full names corresponding to abbreviations are available in **Supplementary Table 5**.

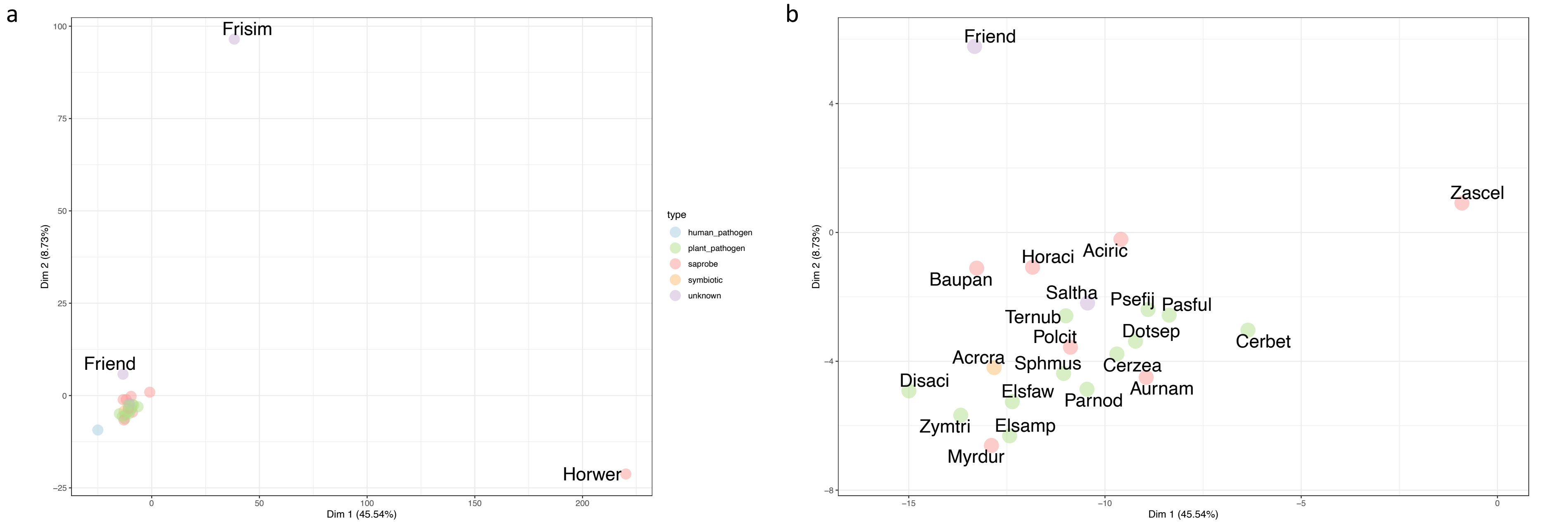

**Supplementary Fig. 14. *Acrodontium* phylogeny with OG losses and gene number.** The blue circles on phylogeny described the loss OG number at each node. The heatmaps showed gene numbers of OGs in representative species. *A. crateriforme* was inferred to loss these OGs.

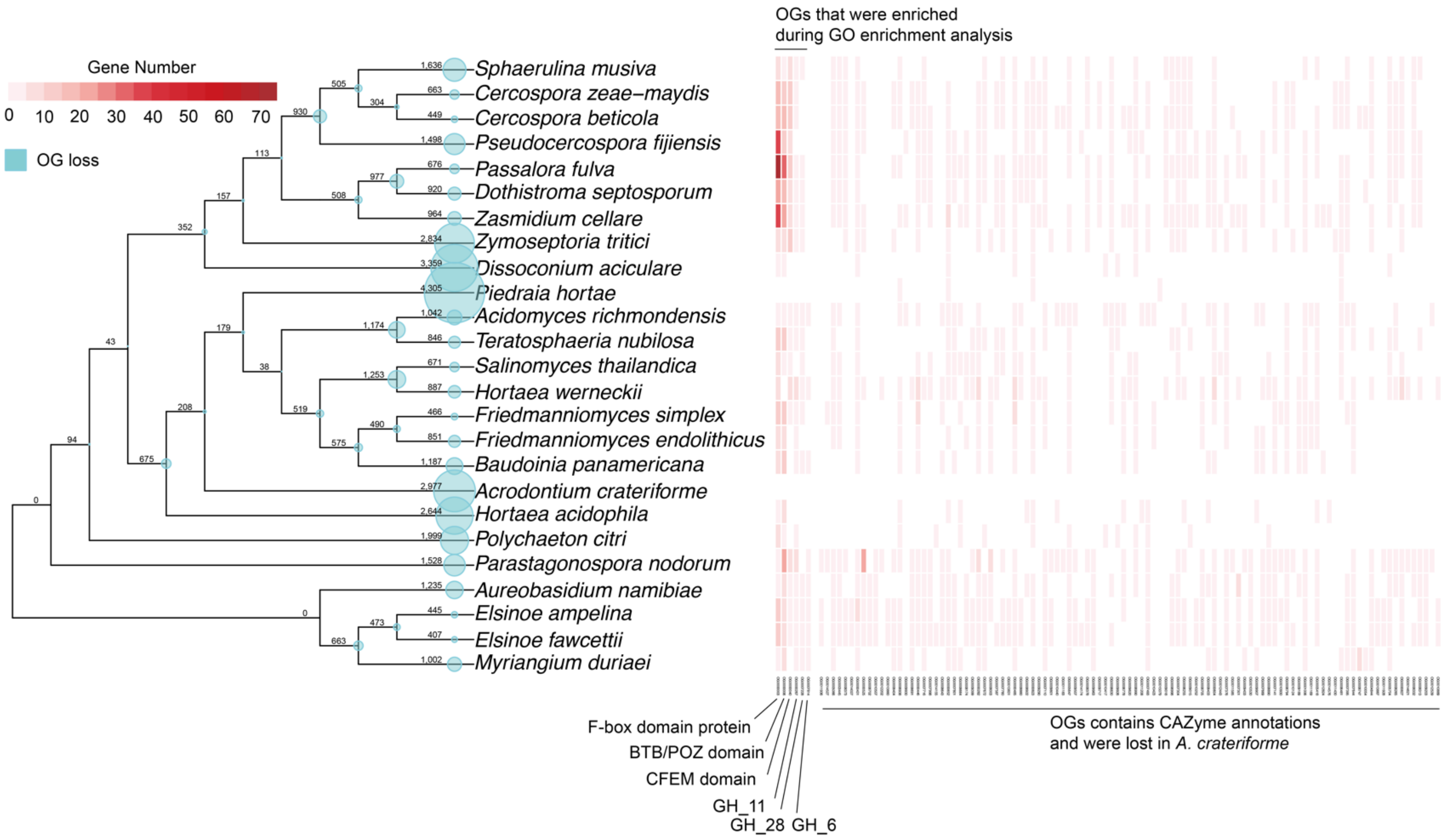

**Supplementary Fig. 15. Top 20 Pfam gain and loss of *A. crateriforme.***
Gains and losses were ranked by domain frequencies. A z-score was calculated for the corresponding abundance of every domain in each species.

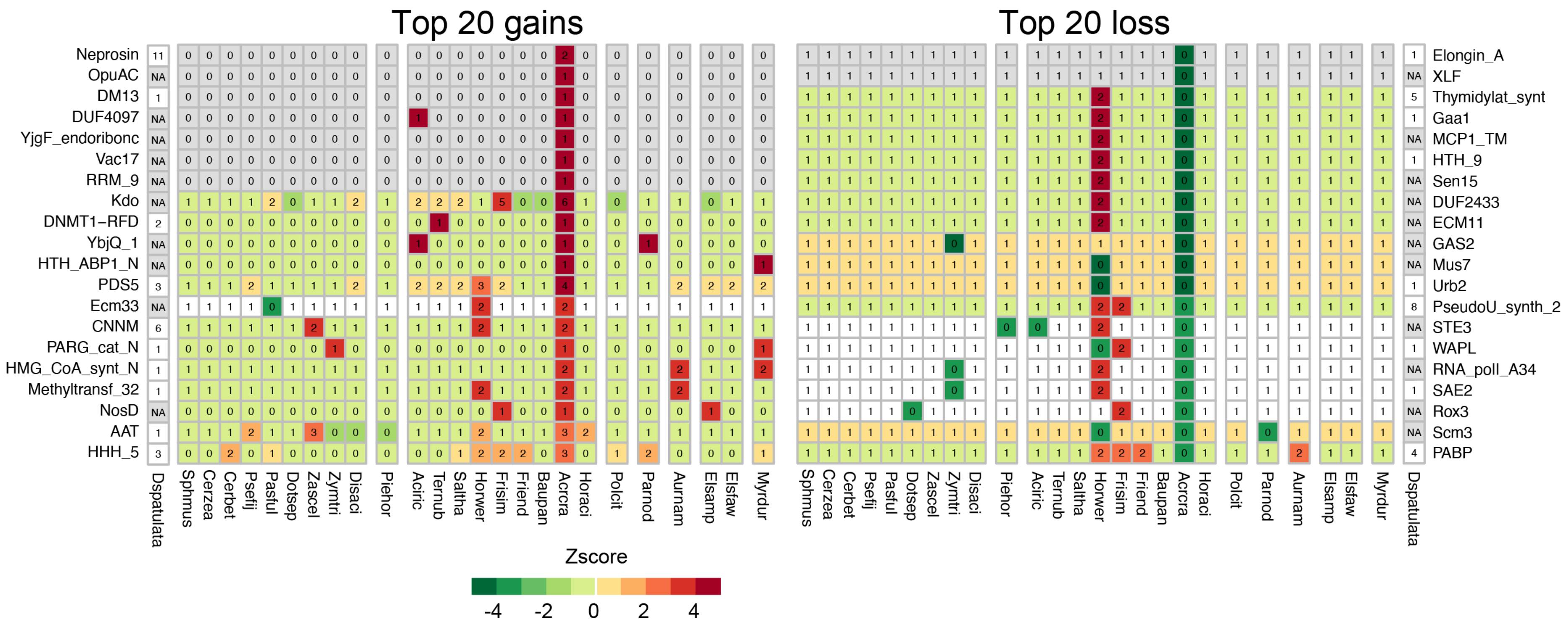

**Supplementary Figure 16. Heatmap of biosynthetic gene clusters (BGC) in 25 fungal species.**

Figure only showed regions with more than 50% orthology to at least one another region.

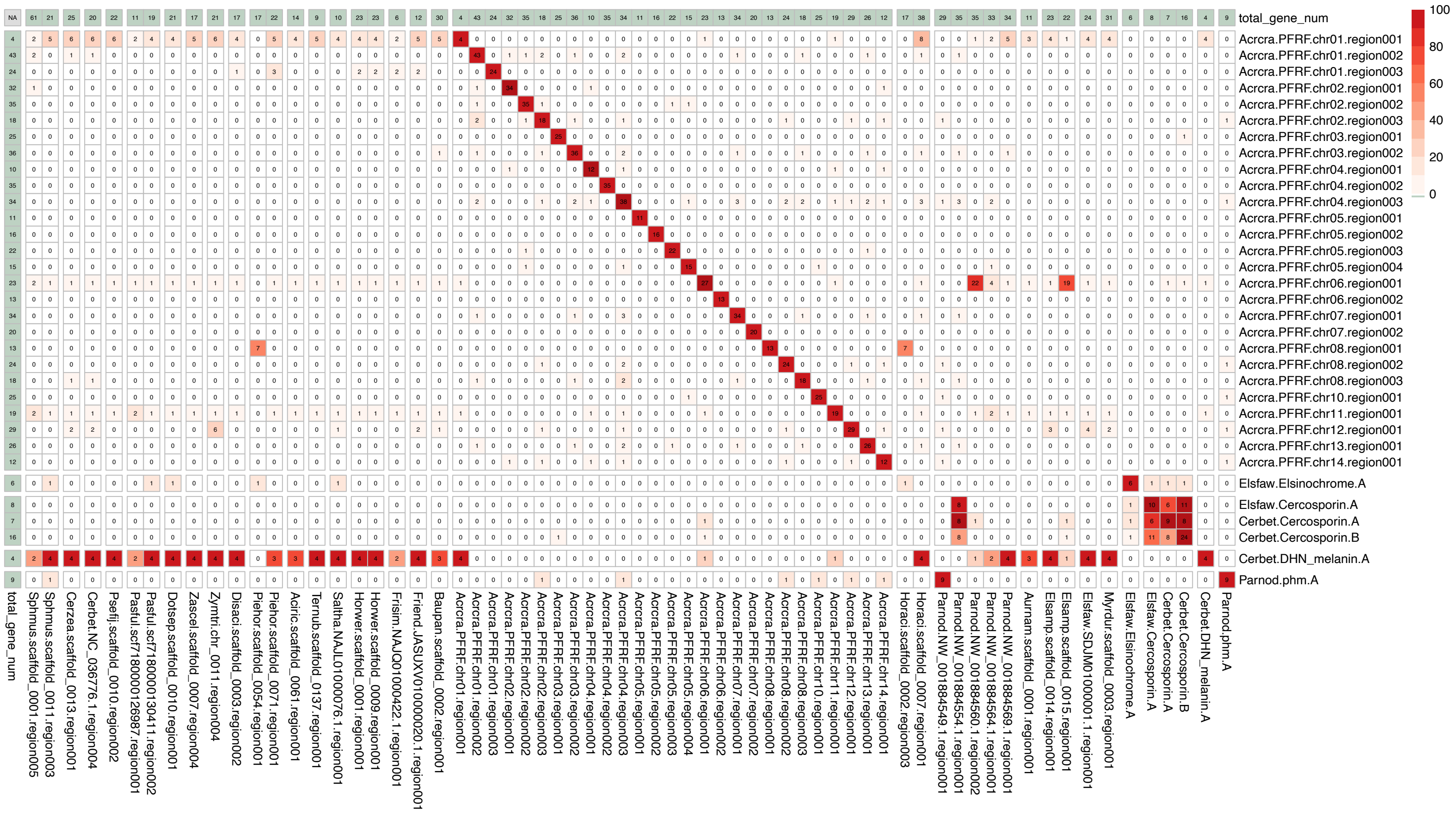

**Supplementary Fig. 17. Gene order within linkage groups has been lost**

Synteny blocks were determined with DAGchainer[39] (ver. r120920) and visualized via CIRCOS[40] (ver. 0.69.9).

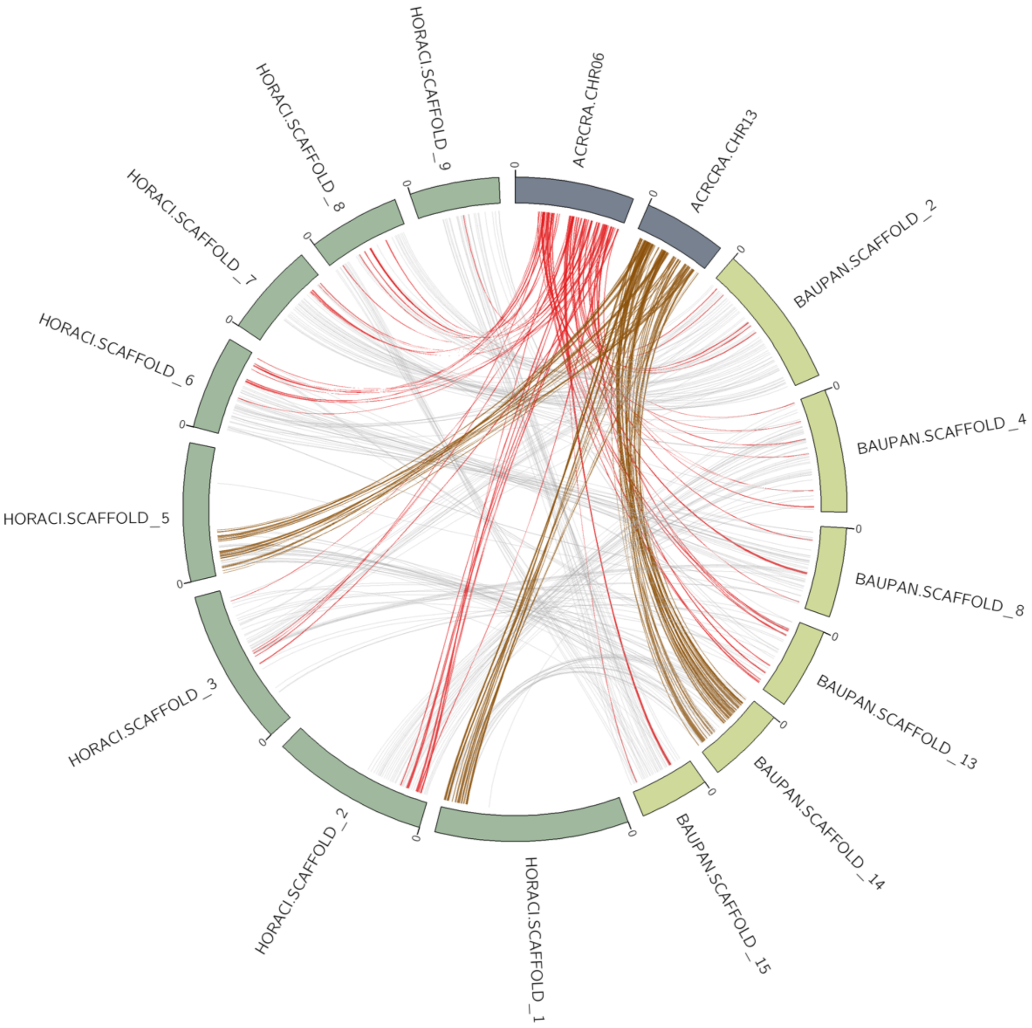

**Supplementary Fig. 18. The location of BGCs in *A. crateriforme*.**

Purple areas are BGC regions, light brown areas show gene density in 10kb sliding windows, and dashed lines show the boundaries of 150kb subtelomere regions.

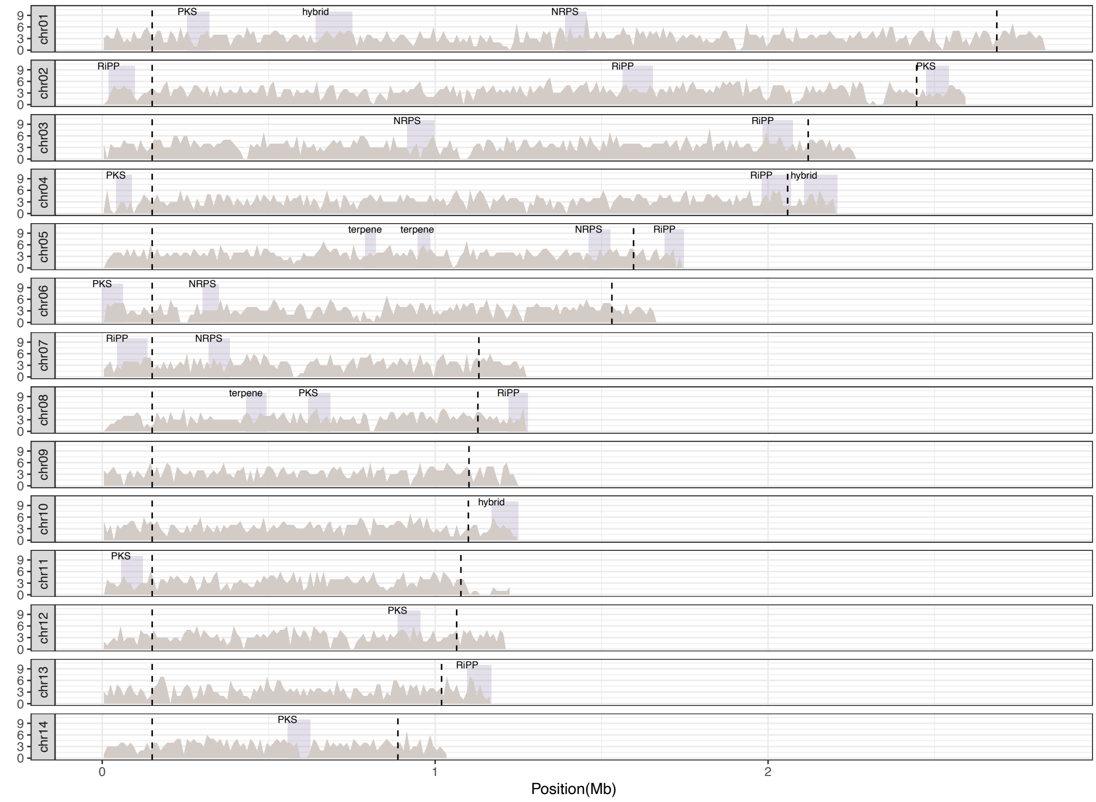

**Supplementary Fig. 19. Schematic diagram of different RNAseq treatment.** Coexistence samples mean *Acrodontium crateriforme* was inoculated on *Drosera spatulata* for one month. Digestion samples show ant powder add on *Drosera spatulata* for 3 days. Coexistence digestion samples show ant powder add on inoculated *Drosera spatulata* for 3 days. To make sure the impact of RNA expression in late stage, we add ant powder add on *Drosera spatulata* for 107 hours.

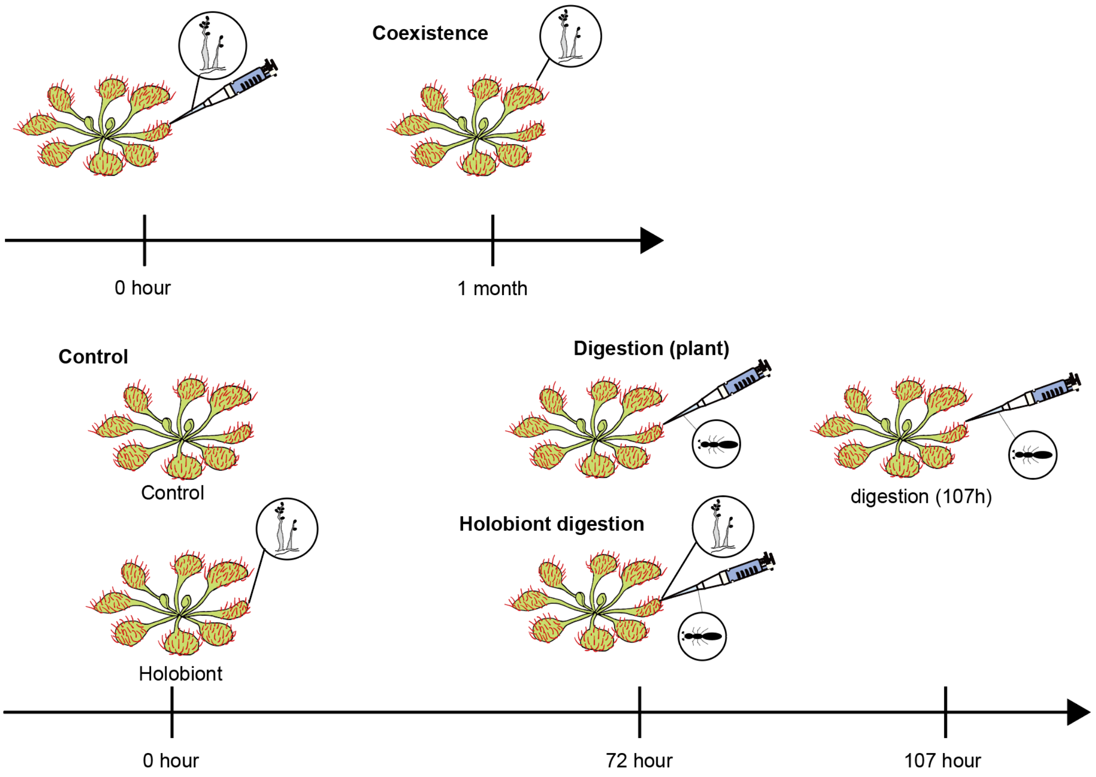

### Supplementary Fig. 20. Overlap of top 20 enriched GO terms in *A. crateriforme* and *D. spatulata* under different conditions.

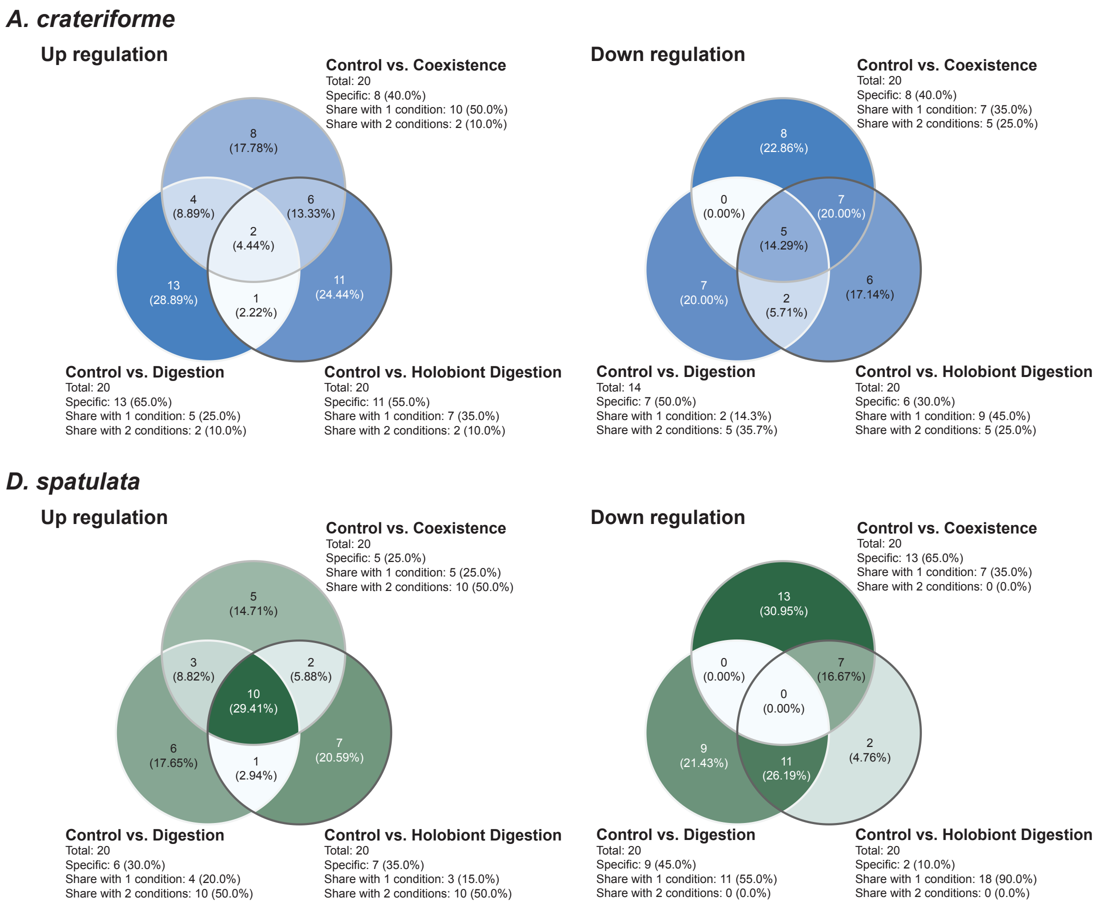

**Supplementary Fig. 21. Expression of sundew chitinase in different treatments**

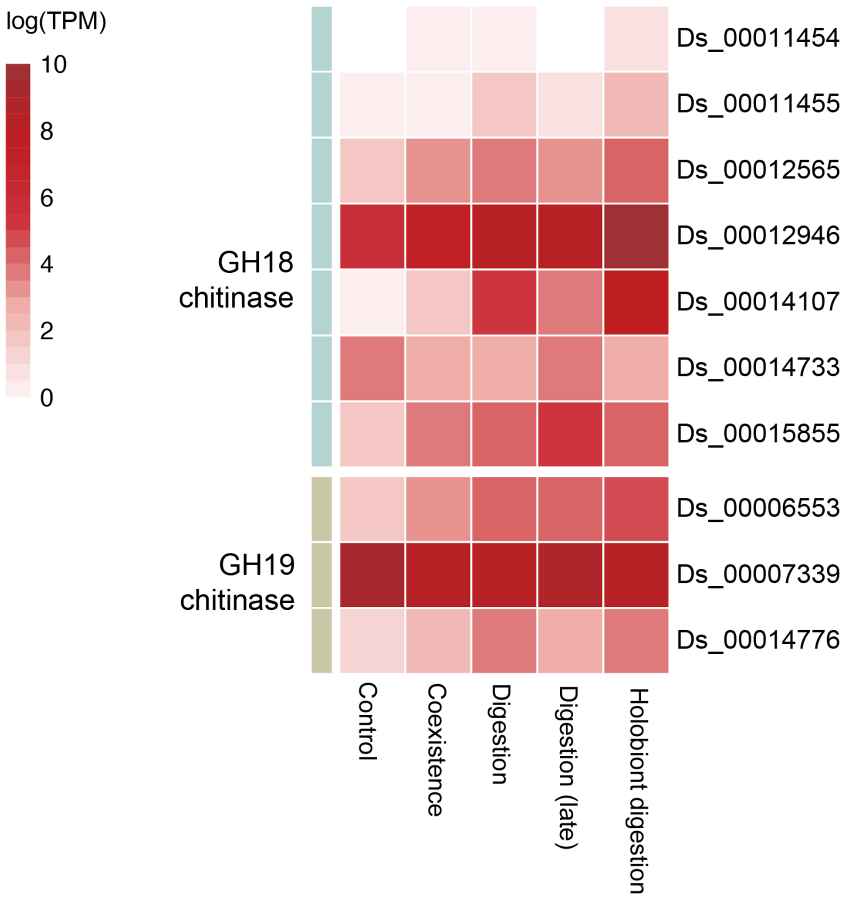

**Supplementary Fig. 22. Expression of ammonium transporters in different treatments**

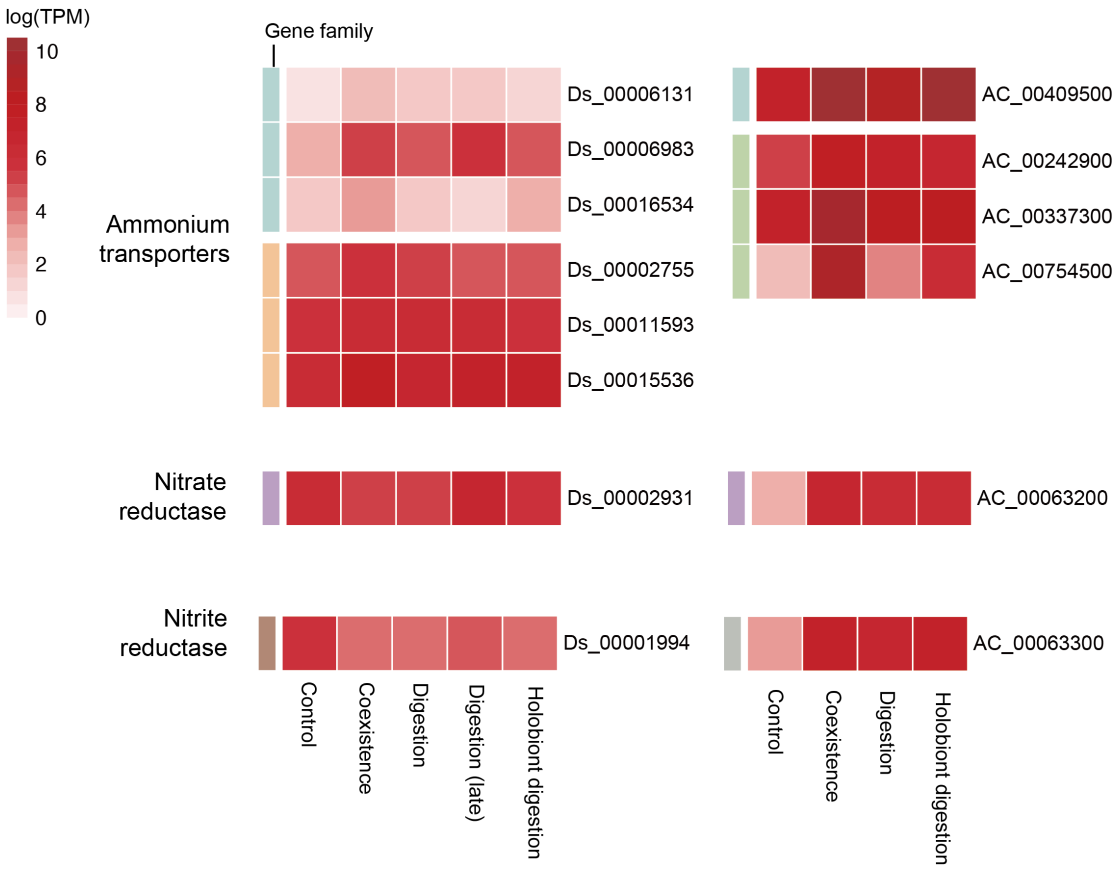

**Supplementary Fig. 23. Co-expression gene modules in *A. crateriforme* across digestion and coexistence conditions using the weighted correlation network analysis (WGCNA)**

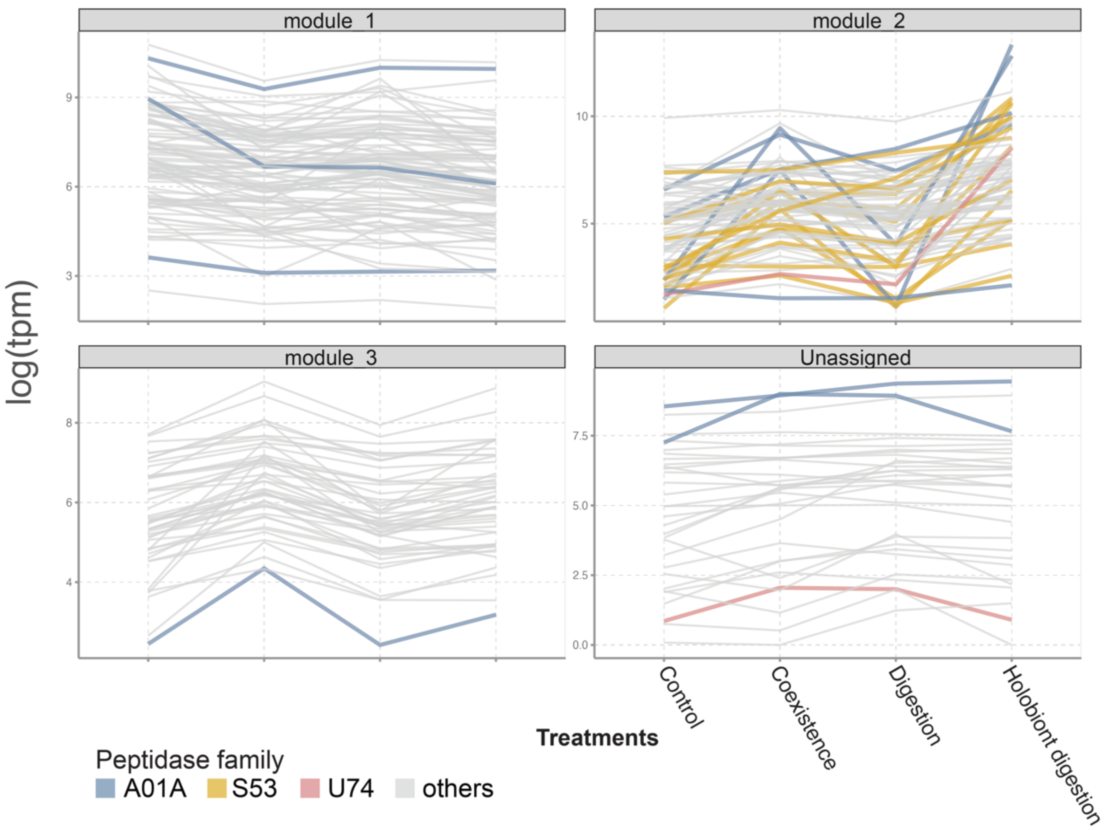

**Supplementary Fig. 24. Co-expression gene modules in *D. spatulata* across digestion and coexistence conditions using the weighted correlation network analysis (WGCNA)**

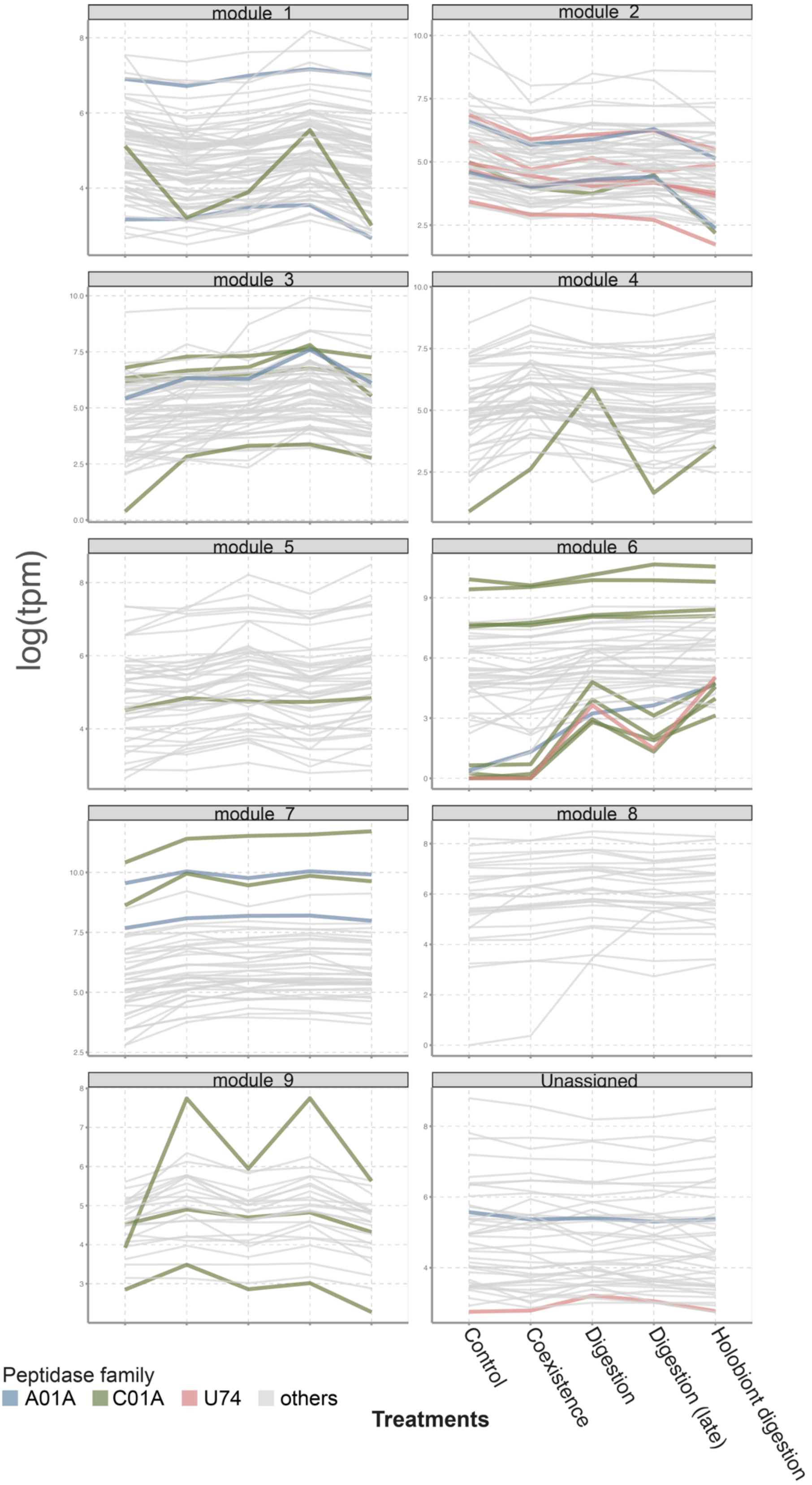

**Supplementary Fig. 25. Expression of fungal transporters in different treatments.**

Star denote significant upregulation in the holobiont digestion phase compared to either digestion or coexistence process.

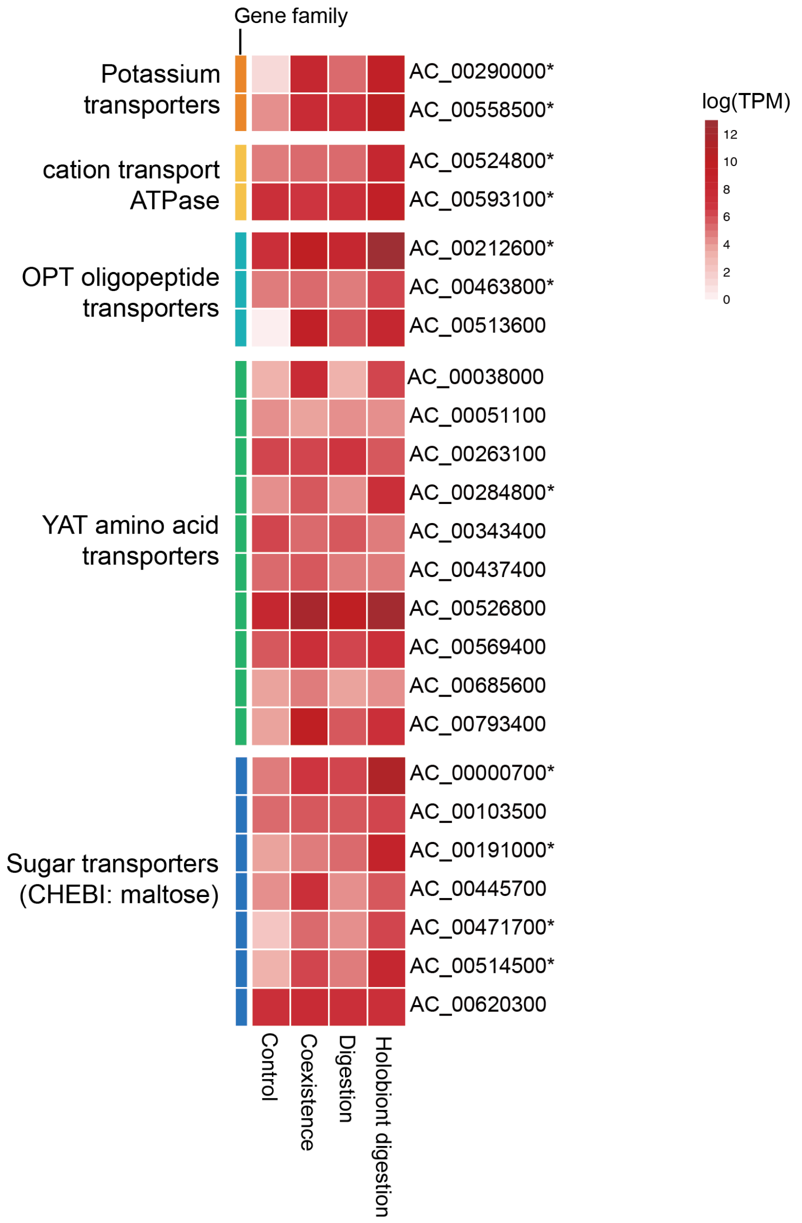

**Supplementary Fig. 26. Phytohormone accumulation in *D. spatulata* in response to different stimuli after two hours.**

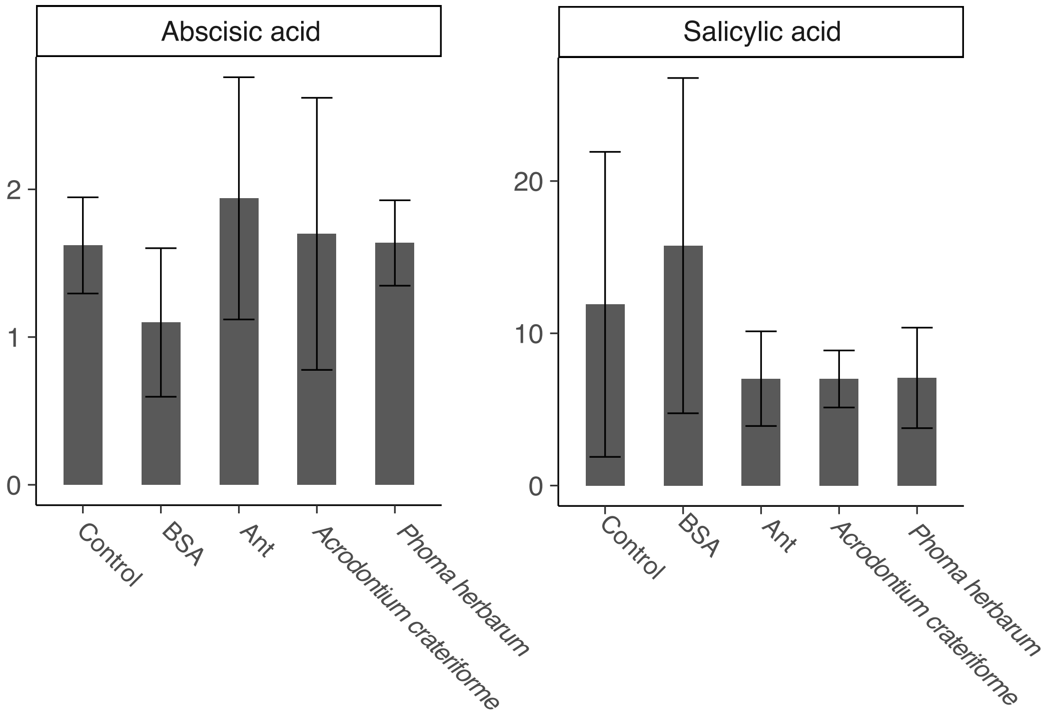

25 Matthew Berriman, A. C., Isheng Jason Tsai Creation of a

comprehensive repeat library for a newly sequenced parasitic worm genome. *PROTOCOL (Version 1) available at Protocol Exchange* doi:https://doi.org/10.1038/protex.2018.054 (2018).
26 Smit, A., Hubley, R & Green, P. RepeatMasker Open-4.0. (2013-2015).
27 Emerson, I. A. & Chitluri, K. K. DCMP: database of cancer mutant protein domains. *Database* **2021**, baab066 (2021).
28 Finn, R. D. *et al.* Pfam: the protein families database. *Nucleic acids research* **42**, D222-D230 (2014).
29 Buchfink, B., Xie, C. & Huson, D. H. Fast and sensitive protein alignment using DIAMOND. *Nature methods* **12**, 59-60 (2015).
30 Elbourne, L. D., Tetu, S. G., Hassan, K. A. & Paulsen, I. T. TransportDB 2.0: a database for exploring membrane transporters in sequenced genomes from all domains of life. *Nucleic acids research* **45**, D320-D324 (2017).
31 Yin, Y. *et al.* dbCAN: a web resource for automated carbohydrate-active enzyme annotation. *Nucleic acids research* **40**, W445-W451 (2012).
32 Rawlings, N. D., Barrett, A. J. & Bateman, A. MEROPS: the peptidase database. *Nucleic acids research* **38**, D227-D233 (2010).
33 Blin, K. *et al.* antiSMASH 6.0: improving cluster detection and comparison capabilities. *Nucleic acids research* **49**, W29-W35 (2021).
34 Huerta-Cepas, J. *et al.* eggNOG 5.0: a hierarchical, functionally and phylogenetically annotated orthology resource based on 5090 organisms and 2502 viruses. *Nucleic acids research* **47**, D309-D314 (2019).
35 Ihaka, R. & Gentleman, R. R: a language for data analysis and graphics. *Journal of computational and graphical statistics* **5**, 299-314 (1996).
36 Alexa, A. & Rahnenführer, J. Gene set enrichment analysis with topGO. *Bioconductor Improv* **27**, 1-26 (2009).
37 Kolde, R. & Kolde, M. R. Package ‘pheatmap’. *R package* **1**, 790 (2015).
38 Wickham, H. ggplot2. *Wiley interdisciplinary reviews: computational statistics* **3**, 180-185 (2011).
39 Haas, B. J., Delcher, A. L., Wortman, J. R. & Salzberg, S. L. DAGchainer: a tool for mining segmental genome duplications and synteny. *Bioinformatics* **20**, 3643-3646 (2004).
40 Krzywinski, M. *et al.* Circos: an information aesthetic for comparative genomics. *Genome research* **19**, 1639-1645 (2009).
